# Dietary tryptophan mitigates lung ischemia-reperfusion injury via microbiota-derived indole-3-propionate and aryl hydrocarbon receptor signaling

**DOI:** 10.64898/2026.04.02.714281

**Authors:** Tomohiro Chaki, Daisuke Maruyama, Thien NM Doan, Tian Xiaoli, Arun Prakash

**Author notes:** Corresponding Author: Arun Prakash, M.D., Ph.D., Department of Anesthesia and Perioperative Care, University of California San Francisco and San Francisco General Hospital, San Francisco, CA. **Abbreviations:** Ischemia-reperfusion, IR; Indole-3-propionate, IPA; IAA; Indole-3-acetate, ILA; Indole-3-lactate, Alveolar macrophage, AM; Aryl-hydrocarbon receptor, AhR.

## Abstract

**Background:** Lung ischemia–reperfusion (IR) injury drives early morbidity after lung transplantation and cardiothoracic surgery, yet targeted preventive therapies are lacking. The gut–lung axis and microbiota-derived tryptophan metabolites, including indole-3-propionate (IPA), may regulate pulmonary immunity and inflammation. We investigated whether a tryptophan-rich (Trp-Rich) diet attenuates sterile lung IR injury by increasing microbiota-derived indole metabolites and reprogramming alveolar macrophage (AM) inflammatory responses.

**Methods:** C57BL/6 mice received isocaloric tryptophan-standard (Trp-Std; 0.18%) or Trp-Rich (0.60%) diets for 14 days, then underwent unilateral left lung IR (60 min ischemia followed by 60 min reperfusion). Oxygen saturation, lung cytokines, and aryl hydrocarbon receptor (AhR) signaling readouts (*Cyp1a1*/*Cyp1b1*) were evaluated. Gut microbiota was profiled by 16S rRNA sequencing, and targeted metabolomics quantified tryptophan metabolites in feces, portal vein (PV) plasma, and lung tissue. To further assess inflammatory priming *in vivo*, mice were additionally challenged with intratracheal lipopolysaccharide (LPS). Mechanistic studies compared IPA with related indole metabolites in MH-S cells and primary human AMs, including *ex vivo* nutritional IR, LPS stimulation, and AhR stimulation and blockade using synthetic agonists and antagonists.

**Results:** Trp-Rich feeding improved post-IR oxygenation, reduced lung IL-1β, and increased pulmonary *Cyp1a1*/*Cyp1b1* gene expression. Trp-Rich diet remodeled the gut microbiota, including enrichment of *Bifidobacterium* and *Lactobacillus*, and increased IPA levels across feces, PV plasma, and lung tissue, with lower kynurenine/IPA ratios across matrices. In the LPS intratracheal challenge, Trp-Rich feeding reduced IL-6 levels in lung tissue and systemic plasma. Primary murine AMs isolated from Trp-Rich mice also showed reduced IL-1β and IL-6 release in an ex vivo nutritional IR model. Among tested indole metabolites, IPA showed the strongest dose-dependent suppression of LPS-induced cytokines and chemokines in MH-S cells and primary human AMs, remained active in the ex vivo nutritional IR model, and its anti-inflammatory effect was abrogated by AhR blockade and enhanced by co-treatment with other indole metabolites.

**Conclusions:** A Trp-Rich diet attenuated sterile lung IR injury, coinciding with gut microbiota remodeling, increased systemic and pulmonary IPA, reduced inflammatory priming, and reprogrammed AM responses. These data support diet- or microbiome-directed strategies targeting IPA–AhR signaling to mitigate perioperative lung IR injury.

**Caption for graphical abstract:** A tryptophan-rich diet remodels the gut microbiota and indole metabolite profiles, including IPA, enhances alveolar macrophage AhR signaling, and attenuates sterile lung ischemia-reperfusion injury.

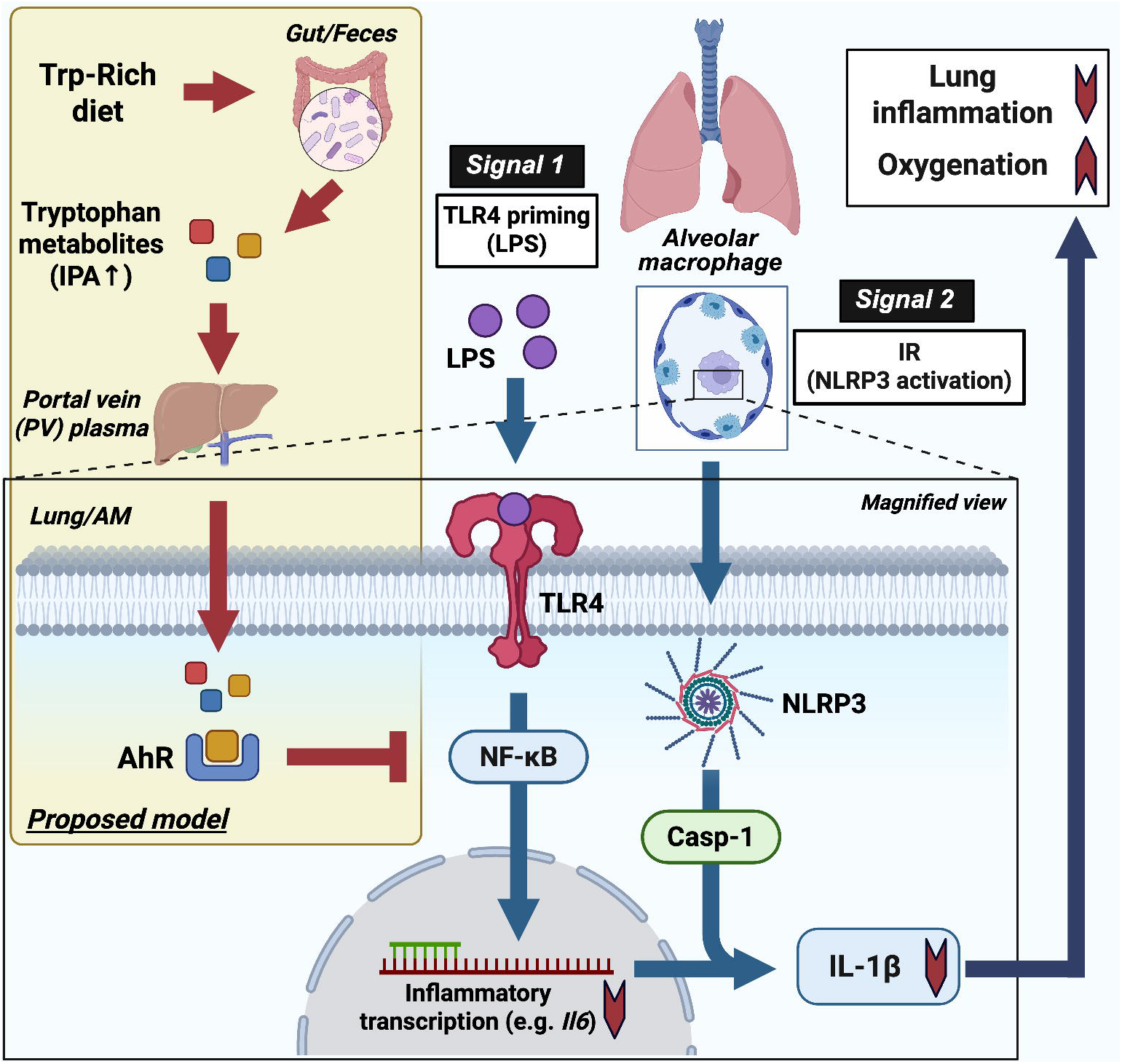

## Introduction

Lung ischemia-reperfusion (IR) injury constitutes a significant contributor to early morbidity and mortality following lung transplantation and cardiothoracic surgical procedures, characterized by an abrupt onset of sterile inflammation, disruption of endothelial and epithelial barriers, and neutrophil-predominant alveolitis [1–3]. Recent experimental and translational investigations have elucidated the pivotal roles of activated alveolar macrophages (AMs) in exacerbating vascular permeability and diffuse alveolar damage [4–8]. Mechanistically, lung IR injury is consistent with a two-signal model in which priming via Lipopolysaccharide (LPS)/TLR4/ NF-κB (Signal 1) and subsequent inflammasome activation triggered by IR-associated cues (Signal 2), including (NOD-like receptor family pyrin domain containing 3 (NLRP3) activation, together drive sterile lung inflammasome [8–11]. Nevertheless, no specific therapeutic interventions for either signal have been incorporated into routine clinical practice. These limitations underscore the need for innovative strategies that modulate upstream inflammatory pathways rather than solely addressing downstream injury. Compelling evidence positions the gut–lung axis as a fundamental regulator of pulmonary immune responses [12]. Alterations in both the composition and functionality of the intestinal microbiota significantly influence susceptibility to pneumonia, acute respiratory distress syndrome (ARDS), and chronic pulmonary diseases through microbially-derived metabolites, such as short-chain fatty acids, tryptophan derivatives, and others, which orchestrate myeloid and epithelial responses at remote mucosal locations (reviewed in [13–18]). Mechanistic investigations reveal that gut-derived metabolites can modulate myeloid and tissue-resident immune cell responses at distant sites [12,19,20]. Conversely, dysbiosis or the depletion of beneficial microbes and metabolites aggravates lung inflammation. These findings suggest that salutary dietary or microbiota-based interventions may provide an indirect protective effect against lung IR injury by reprogramming systemic and pulmonary immune responses [21]. Nonetheless, this concept remains inadequately explored within the context of acute sterile lung injury.

Among microbial metabolites, indole derivatives generated from dietary tryptophan are emerging as key regulators of mucosal immunity (reviewed in [22]). Indole-3-propionate (IPA) is a prominent member of the microbiota-derived indole family, which also includes indole-3-acetate (IAA) and indole-3-lactate (ILA). IPA exerts antioxidant and anti-inflammatory effects across experimental models, including suppression of NF-κB and inflammasome signaling, preservation of barrier integrity, and modulation of macrophage phenotypes via aryl hydrocarbon receptor (AhR) signaling [23–25]. Microbiota-derived indoles can act through AhR-dependent pathways to regulate immune responses at barrier and distant tissues [19,20]. Microbiota-derived indoles have also been linked to protection against systemic inflammation and mitigation of mucosal and pulmonary injury in infection-related models [26,27]. However, it remains unclear whether a dietary strategy that increases microbial indole metabolism can raise endogenous IPA across the gut and circulation, particularly within the lung, and whether these changes are coupled with reduced lung inflammation and altered AhR-related responses during sterile lung IR injury.

Consequently, we posited that a diet rich in tryptophan would modify gut microbiota composition and enhance the production of gut-derived tryptophan metabolites, particularly IPA, reducing lung inflammation following IR injury. Utilizing a murine lung IR model in conjunction with targeted metabolomics, gut microbiota profiling, and *in vitro* assays with MH-S and human AMs, we sought to elucidate the gut–lung axis that underlies the protective effects mediated by tryptophan metabolites in acute lung IR injury.

## Materials and Methods

Methods are summarized here. Full experimental procedures and analysis parameters are provided in the Supplemental Methods.

### Animal care

All animal procedures were approved by the Institutional Animal Care and Use Committee at the University of California San Francisco (protocol: AN197325-01). Wild-type C57BL/6 mice (10-15 weeks old, both sexes) were housed under standard conditions with free access to food and water. Mice were randomized to an isocaloric, isonitrogenous tryptophan-standard diet (Trp-Std, 0.18%) or tryptophan-rich diet (Trp-Rich, 0.60%) for 14 days (Inotiv). Group sizes were Trp-Std (males n = 14, females n = 13) and Trp-Rich (males n = 12, females n = 11) in the lung IR experiment, Trp-Std (males n = 10) and Trp-Rich (males n = 8) in the intratracheal LPS challenge experiment, and Trp-Std (males n = 10) and Trp-Rich (males n = 10) in the primary AMs experiment. Cellulose content was matched between diets (3%) to minimize fiber-driven microbiome differences (Supplemental Table S1). Of note, body weight did not differ between groups (Figure S1A), and gross intestinal appearance was comparable after 2 weeks of dietary intervention (Figure S1B).

### Left lung IR surgery and sample collection

Unilateral left lung IR was performed as previously described [8,28,29]. In brief, mice were anesthetized, intubated, and mechanically ventilated, and the left pulmonary artery was occluded using a reversible slipknot for 60 min, followed by 60 min of reperfusion. Peripheral oxygen saturation and respiratory rate were measured immediately prior to euthanasia (MouseOx Plus) and assessed in male mice only, as the MouseOx neck-collar sensor is recommended for animals >25 g, and its use in smaller female mice could compromise breathing. Blood was collected from the inferior vena cava and portal vein (PV) into heparinized syringes, centrifuged, and the plasma was stored at -80°C. The left lung was divided for RNA extraction and cytokine quantification, and the right lung was used for targeted metabolomics. Fecal pellets were collected from the colon and stored at -80°C for 16S rRNA sequencing and metabolomics.

### Intratracheal LPS challenge

To assess LPS/TLR4-driven inflammatory priming (Signal 1) *in vivo*, a subset of mice after 2 weeks of dietary intervention received intratracheal LPS (5 mg/kg). Lung tissue and inferior vena cava (IVC) plasma were collected 6 h later for cytokine quantification.

### Reagents and Cell Lines

IPA, ILA, IAA, indole, L-tryptophan, LPS (*E. coli* 0111:B4), and the AhR antagonist CH-223191 were purchased from Sigma-Aldrich. TCDD was purchased from Neta Scientific. MH-S cells (murine AM line) were obtained from ATCC and cultured in RPMI 1640 supplemented with 10% fetal bovine serum (FBS) and 1% penicillin-streptomycin.

### Collection of Human Primary AMs

Primary human AMs were obtained by bronchoalveolar lavage from adult donor lungs declined for transplantation (UCSF, provided by Dr. Michael Matthay). Cells were processed to remove erythrocytes, resuspended in RPMI 1640 with 10% FBS and antibiotics, and allowed to adhere before stimulation experiments. Donor information is summarized in Supplemental Table S2.

### *In vitro* and *Ex vivo* AM assays

For MH-S experiments, cells were pretreated with tryptophan-related metabolites (IPA, ILA, IAA, indole, or L-tryptophan; 0.01-1 mM) and then co-treated with LPS (50 ng/mL) in the continued presence of the metabolites. For AhR-dependence experiments, cells were pretreated with CH-223191 and then exposed to IPA (0.01-0.1 mM) or TCDD (10 nM) followed by LPS co-treatment, with or without CH-223191. For Human AM experiments, cells were co-treated with LPS (10 ng/mL) and IPA (0.01-1 mM) overnight. *Ex vivo* nutritional IR using human and mouse primary AMs was performed as previously described [21,28]. Briefly, we co-treated primary AMs with LPS (200 ng/mL) and IPA overnight, washed, incubated in phosphate-buffered saline (PBS) for 1 h (ischemia-mimicking phase), and then switched to RPMI 1640 medium (reperfusion-mimicking phase). Supernatants were collected for cytokine measurements, and in the human AM experiment, cell number and confluency were quantified by imaging for normalization.

### Cytokine measurement and gene expression

Mouse lung homogenates and cell-culture supernatants were analyzed using DuoSet ELISA kits (R&D Systems). Total RNA was extracted from lung tissue or cells, reverse-transcribed, and analyzed by RT-qPCR using TaqMan assays. Relative gene expression was calculated using the 2^-ΔΔCt method with *Actb*, *Gapdh*, and *Rpl19* as reference genes.

### 16S rRNA gene sequencing and microbiome analysis

For unbiased microbiome analysis, we analyzed the first four mice enrolled per group (n = 4 per group). Microbial DNA was extracted from fecal samples (QIAamp PowerFecal Pro DNA Kit, Qiagen). The V4 region of the bacterial 16S rRNA gene was amplified using the 515F/806R primer pair, pooled, purified, and sequenced on an Illumina MiSeq platform (2×300 bp). Demultiplexed paired-end reads were subsequently processed using a DADA2-based workflow in MicrobiomeAnalyst, and taxonomy was assigned against SILVA v138 [30,31].

### Targeted metabolomics of tryptophan-related metabolites

Tryptophan-related metabolites were quantified in feces, PV plasma, and lung using targeted LC-MS with isotope-labeled internal standards and external calibration performed by the metabolomics core at the University of Michigan (Maureen Kachmann). Samples were extracted using matrix-specific solvent protocols, analyzed with pooled quality-control injections, and quantified using MassHunter software. Tissue concentrations were normalized to input mass.

### Multi-omics integration and statistical analysis

Microbiome profiles were generated in MicrobiomeAnalyst using a DADA2-based ASV workflow with standard feature filtering [30,31]. β-diversity was evaluated using Bray-Curtis and UniFrac distances and tested by PERMANOVA. Differential abundance was assessed using DESeq2 with FDR correction, and predicted functional potential was inferred using Tax4Fun2 (KEGG ortholog profiles). Targeted metabolomics data were analyzed in MetaboAnalyst 6.0 with log transformation and scaling, with group comparisons performed using FDR-adjusted univariate testing and visualization by ordination and volcano plots [32,33]. Associations between taxa, metabolites, and lung inflammatory/AhR readouts were assessed using Spearman correlations. All other statistical analyses were performed in GraphPad Prism 10 (GraphPad Software, Boston, MA). Data are presented as mean ± SD. Two-group comparisons used unpaired t-tests, or Mann-Whitney tests as appropriate. Multi-group comparisons were performed using one-way ANOVA with Tukey’s post hoc test. A two-sided p-value < 0.05 was considered significant.

## Results

### A Trp-Rich diet attenuates lung IR injury and activates AhR signaling

To assess the *in vivo* impact of dietary tryptophan, mice were fed a Trp-Rich or Trp-Std diet for 2 weeks (Figure 1A). After left lung IR, oxygen saturation was higher in the Trp-Rich group despite similar respiratory rates, indicating improved lung function (Figures 1B-C). In left lung homogenates, IL-1β was reduced in the Trp-Rich group, whereas IL-6 did not differ between groups (Figures 1D-E). Given prior evidence that estrogen can mitigate lung IR injury [34], we examined sex-specific responses. The protective phenotype was not observed in females but was preserved in males (Figures S1C-D). Therefore, subsequent analyses were performed in male mice, and the sex-dependent effects of dietary tryptophan on lung injury responses warrant further study, in keeping with prior reports that lung IR injury is modulated by sex and estrogen signaling [34]. Lung expression of the canonical AhR target genes *Cyp1a1* and *Cyp1b1* was increased in the Trp-Rich group, indicating successful enhancement of AhR signaling (Figures 1F-G) [35,36].

**Figure 1.**
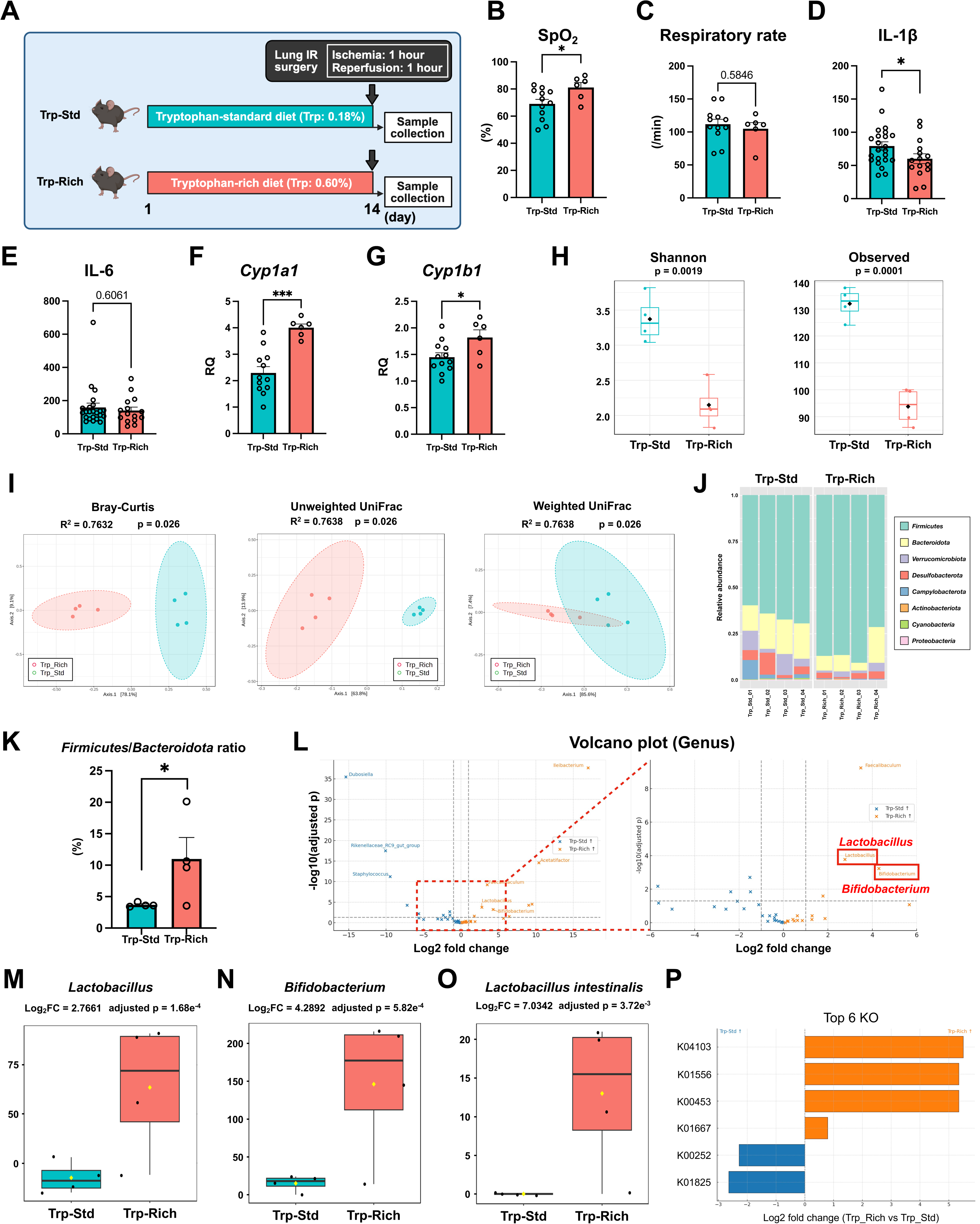
Tryptophan-rich diet attenuates lung ischemia-reperfusion injury and remodels the gut microbiota. **A)** Study design. Mice were assigned to isocaloric tryptophan-standard (Trp-Std, 0.18% tryptophan) or tryptophan-rich (Trp-Rich, 0.60% tryptophan) diets for 2 weeks, followed by unilateral lung ischemia-reperfusion (60 min ischemia and 60 min reperfusion) and sample collection. **B-C)** Oxygen saturation (SpO_2_) and respiratory rate measured after 60 min of reperfusion. SpO_2_ was higher in Trp-Rich mice, whereas respiratory rate did not differ between groups (Trp-Std, n = 12; Trp-Rich, n = 6). **D-E)** Lung inflammatory cytokines quantified by ELISA showing reduced IL-1β in the Trp-Rich group, with no significant difference in IL-6 (Trp-Std, n = 23; Trp-Rich, n = 15). **F-G)** RT-qPCR analysis of aryl hydrocarbon receptor (AhR) target genes (*Cyp1a1* and *Cyp1b1*) demonstrating increased expression in Trp-Rich lungs (Trp-Std, n = 12; Trp-Rich, n = 6). **H)** Alpha-diversity metrics (Shannon index and observed features) showing reduced diversity in the Trp-Rich group. **I)** β-diversity ordinations based on Bray-Curtis, unweighted, and weighted UniFrac distances demonstrating separation of Trp-Std and Trp-Rich communities (PERMANOVA). **J)** *Phylum* level relative abundance and **K)** *Firmicutes*/*Bacteroidota* ratio, which was increased by the Trp-Rich diet. **L)** *Genus* level differential abundance volcano plot highlighting taxa altered by diet. **M-O)** Relative abundances of *Lactobacillus*, *Bifidobacterium*, and *Lactobacillus intestinalis*, each enriched in Trp-Rich mice. The taxonomy chart at the *genus* level and volcano plot at the *species* level are presented in Supplemental Figure S2. **L)** Predicted functional profiling of the tryptophan metabolism pathway (ko00380) showing enrichment of predicted KEGG orthologs related to indole-kynurenine metabolism in the Trp-Rich group (K04103, K01556, K00453, and K01667). PERMANOVA was used for β-diversity comparisons, and other comparisons were performed using an unpaired t-test or the Mann-Whitney test with false discovery rate correction for multiple testing. n = 4 in each group in microbiome analysis. *: p<0.05, ***: p<0.001.

### Remodeling of gut microbiota by a Trp-Rich diet

Using fecal 16S rRNA sequencing, we evaluated how a Trp-Rich diet alters gut microbial communities. Alpha-diversity was lower in the Trp-Rich group compared with the Trp-Std group (Figures 1H and S2A-D). Beta-diversity analyses also demonstrated clear group separation across Bray-Curtis, unweighted UniFrac, and weighted UniFrac distances (Figure 1I). At the *phylum* level, the Trp-Rich diet increased *Firmicutes* and decreased *Campylobacterota* and *Cyanobacteria*, resulting in a higher *Firmicutes*/*Bacteroidota* ratio (Figures 1J-K, Supplemental Table S3). At the *genus* level, *Faecalibaculum* and *Dubosiella* were dominant taxa (Figure S2E). Differential abundance analysis identified several taxa enriched in the Trp-Rich group at the *genus* level, including *Lactobacillus* and *Bifidobacterium*, whereas *Dubosiella*, *Rikenellaceae_RC9_gut_group*, and *Staphylococcus* were relatively depleted (Figure 1L, Supplemental Table S4). At the putative *species* level based on 16S V4 taxonomic assignment, *Lactobacillus intestinalis*, *Ileibacterium valens*, and *Faecalibaculum rodentium* were enriched in the Trp-Rich group, while *Streptococcus danieliae and Helicobacter ganmani* were depleted (Figure S2F, Supplemental Table S5). The expansion of *Lactobacillus*, *Bifidobacterium*, and *Lactobacillus intestinalis* in Trp-Rich mice (Figures 1M-O) was observed, and these taxa have been linked to increased production of indole-related tryptophan metabolites, including ILA and IPA, in the gut [37,38]. Consistent with these taxonomic shifts, predicted functional profiling (Tax4Fun2, SILVA) followed by metagenomeSeq analysis indicated higher predicted abundance of key KEGG orthologs in the tryptophan metabolism pathway in the Trp-Rich group, including K04103 (ipdC), K01667 (tnaA), and K00453 (TDO/TDO2) within the tryptophan metabolism pathway (ko00380), supporting increased community-level potential for indole-branch metabolism consistent with downstream IPA and other indole derivatives production (Figure 1P).

### Transmission of tryptophan metabolites along the gut-lung axis with a Trp-Rich diet

Targeted metabolomics of tryptophan-related metabolites in feces, PV plasma, and right lung were performed to assess how a Trp-Rich diet reshapes tryptophan metabolic profiles. Principal component analysis demonstrated distinct metabolic profiles between Trp-Std and Trp-Rich groups across all matrices (Figure 2A). Volcano plots showed that IPA was consistently increased in the Trp-Rich group in feces, PV plasma, and lung (Figures 2B-E). In addition, kynurenic acid and 5-hydroxyindoleacetic acid were increased in feces in the Trp-Rich group, whereas kynurenine in the lung was increased in the Trp-Std group. To capture pathway-level balance, we calculated kynurenine/IPA and kynurenine/tryptophan ratios [39,40]. The kynurenine/IPA ratio was lower in feces, PV plasma, and lung in the Trp-Rich group (Figure 2F). The kynurenine/tryptophan ratio was also reduced in PV plasma, with no differences in feces or lung (Figure 2G). In contrast, ILA and IAA did not differ between groups in any matrix (Figure S3). Together, these data indicate that the Trp-Rich diet shifted tryptophan metabolism toward the indole branch, characterized by increased IPA relative to kynurenine.

**Figure 2.**
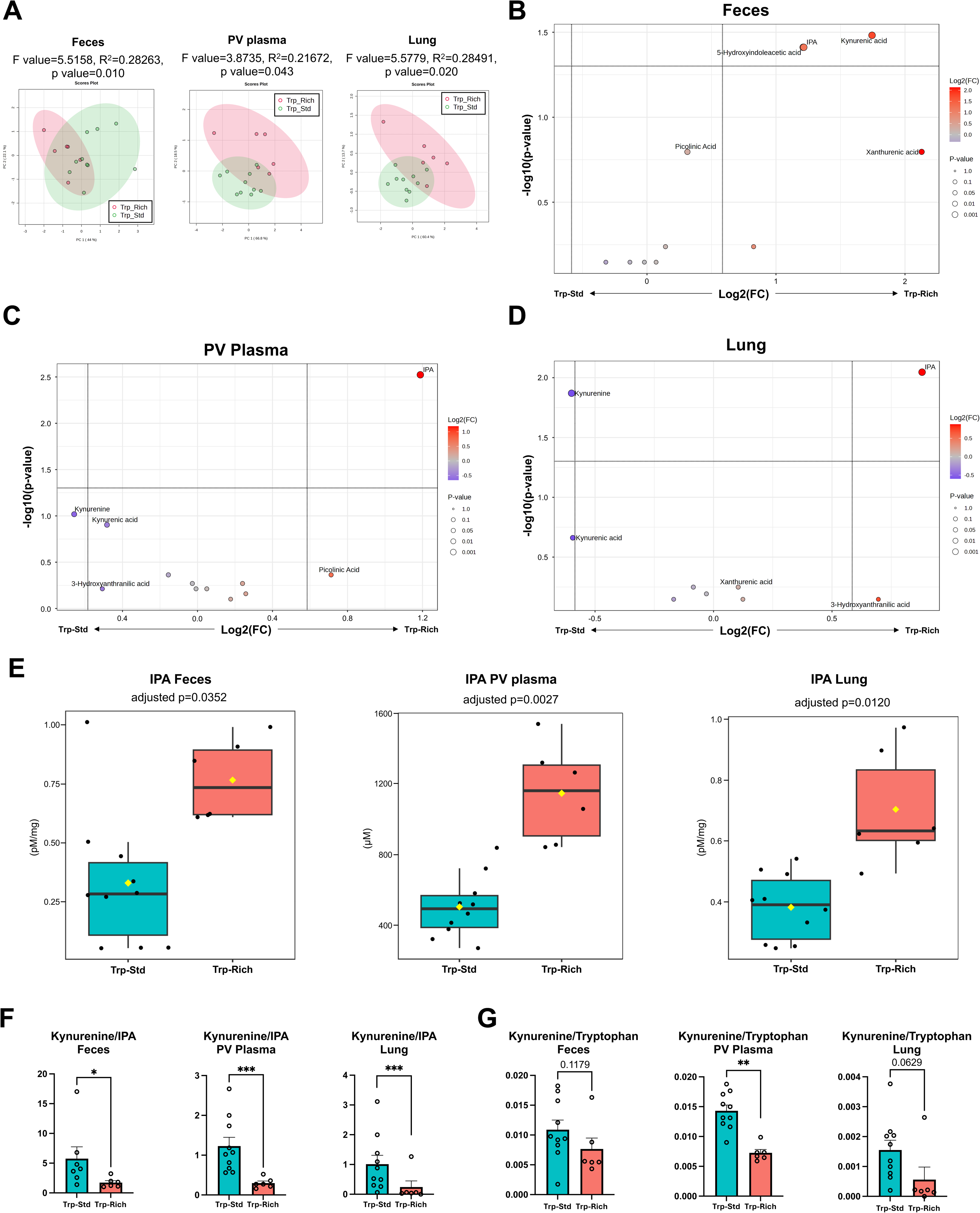
A tryptophan-rich diet reshapes tryptophan metabolism. **A)** Principal component analysis (PCA) of targeted tryptophan metabolite profiles in feces, portal vein (PV) plasma, and lung tissue, showing separation of tryptophan-standard (Trp-Std) and Trp-Rich groups in each matrix. **B-D)** Volcano plot demonstrating differential abundance of targeted tryptophan metabolites in feces, PV plasma, and lung tissue between Trp-Std and Trp-Rich groups. The complete panels of targeted tryptophan metabolites are shown in Supplemental Figures S3. **E)** Indole-3-propionate (IPA) concentrations in the feces, PV plasma, and lung, demonstrating increased IPA in Trp-Rich mice across all three matrices. **F)** Kynurenine/IPA ratios in feces, PV plasma, and lung, showing reduced ratios in the Trp-Rich group. **G)** Kynurenine/tryptophan ratios in feces, PV plasma, and lung, with a significant reduction in PV plasma in Trp-Rich mice. Together, these ratio-based metrics support a relative shift in pathway balance toward indole/IPA production with Trp-Rich feeding as indirect surrogates rather than direct flux measurements. PERMANOVA evaluated group separation in PCA space, and other comparisons were performed using an unpaired t-test or the Mann-Whitney test with false discovery rate correction for multiple testing. Trp-Std, n = 10; Trp-Rich, n = 6. *: p<0.05, **: p<0.01, ***: p<0.001.

### Integrated multi-omics analyses

To link diet-induced changes across the microbiome, metabolome, and host lung readouts, multi-omics correlation analyses were performed [30–32,41]. At the *genus* level, *Bifidobacterium* was inversely correlated with lung IL-1β and IL-6 (Figures 3A and S4A-B), whereas correlations between *Lactobacillus* and these cytokines were not significant (Figures 3A, S4A, and S4C). At the *species* level, *Lactobacillus intestinalis* showed an inverse correlation with lung IL-6 (Figures S4D-E). Vis-à-vis AhR signaling, *Lactobacillus intestinalis* was positively correlated with lung *Cyp1b1* (Figures 3B and S4E), however correlations involving *Bifidobacterium* or *Lactobacillus* with *Cyp1a1* or *Cyp1b1* were not significant (Figure S4B-C).

**Figure 3.**
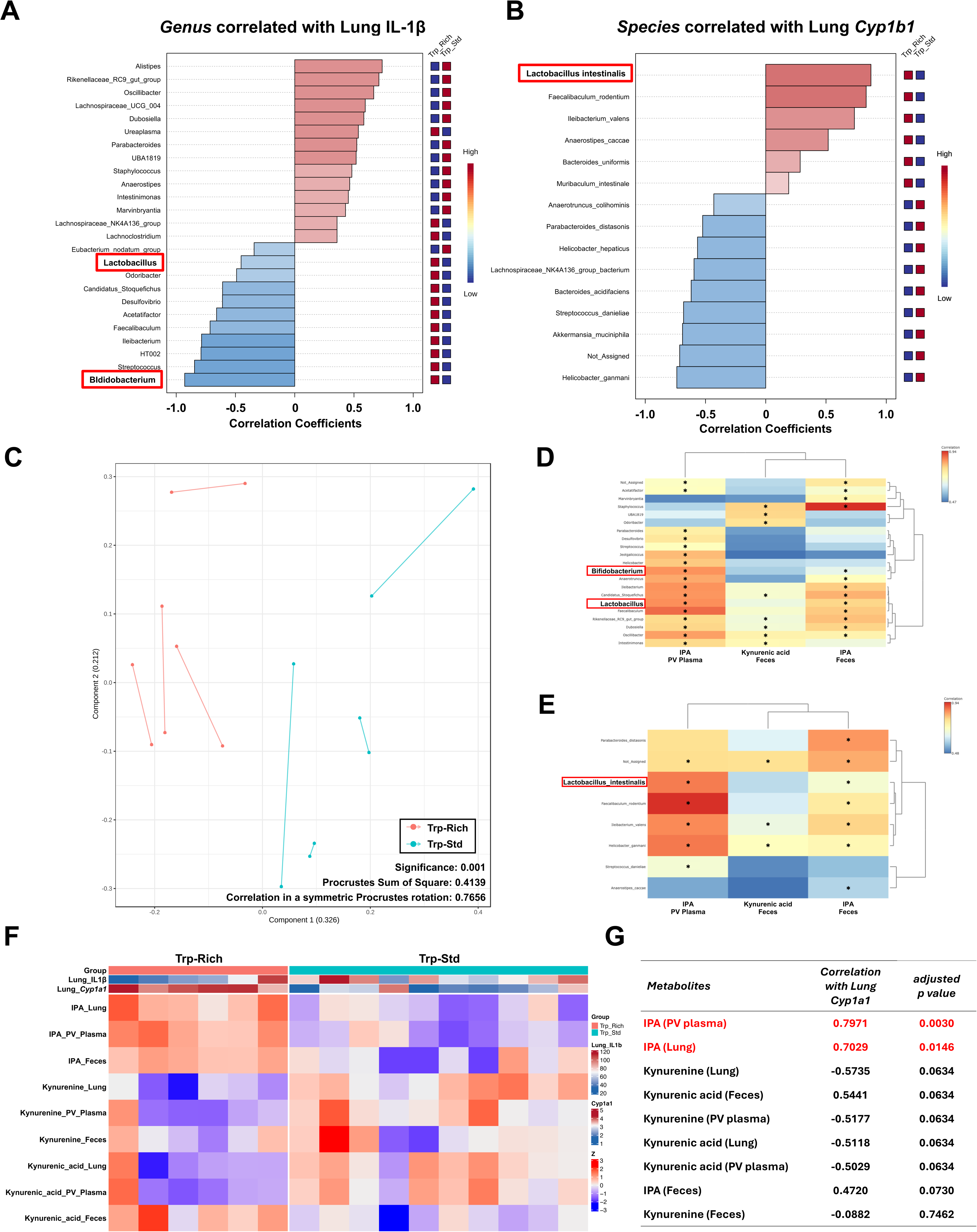
Integrated multi-omics analyses among gut microbiome, tryptophan metabolism, and lung inflammatory/aryl hydrocarbon receptor signaling readouts. **A)** *Genus* level correlations between gut microbiota and lung inflammatory markers showing *Bifidobacterium* abundance was inversely correlated with lung IL-1β. **B)** *Species* level correlations showing that *Lactobacillus intestinalis* abundance was positively correlated with lung *Cyp1b1* expression. Additional correlation analyses are provided in Supplemental Figure S4. C) Procrustes analysis comparing microbiome and metabolomics ordinations, indicating overall concordance between datasets based on generally short Procrustes distances. **D-E)** Correlation screening/heatmap analysis linking gut taxa with key metabolites, highlighting positive associations of *Bifidobacterium*, *Lactobacillus*, and *Lactobacillus intestinalis* with indole-3-propionate (IPA) in PV plasma. **F-G)** Correlation heatmap/plots linking lung *Cyp1a1* expression with IPA levels in PV plasma and lung. Associations were assessed using Spearman’s rank correlation with false discovery rate correction. Trp-Std, n = 10; Trp-Rich, n = 6. In microbiome-related analysis, n = 4 in each group. *: p<0.05.

Procrustes analysis indicated overall concordance between microbiome and metabolomics ordinations, with a subset of samples showing larger discrepancies (Figure 3C). Correlation analysis further identified significant associations between *Bifidobacterium* and *Lactobacillus* and PV plasma IPA at the *genus* level (Figures 3D and S4F), and between *Lactobacillus intestinalis* and PV plasma IPA at the *species* level (Figures 3E and S4F). Finally, PV plasma and lung IPA levels were positively correlated with lung *Cyp1a1* (Figures 3F-G and S4G-H). Collectively, these integrated analyses support a model in which a Trp-Rich diet remodels the gut microbiota and tryptophan metabolism, increasing key taxa and IPA, which are linked to AhR-related responses and reduced inflammatory lung phenotype post-IR injury.

### A Trp-Rich diet suppresses LPS-induced IL-6 responses *in vivo*

To specifically test whether a Trp-Rich diet modulates inflammatory priming (Signal 1), we performed an intratracheal LPS challenge (5 mg/kg) and quantified IL-6 in lung tissue and IVC plasma at 6 h (Figure 4A). Lung and plasma IL-6 levels were significantly lower in Trp-Rich mice, indicating attenuated pulmonary and systemic cytokine responses to LPS (Figures 4B-C). Consistent with reduced IL-6 protein, lung *Il6* transcript levels were also lower at 6 h after IT LPS challenge (Figure S5A). In contrast, lung IL-1β protein and gene expression and TNF-α levels did not differ significantly between groups (Figure S5B-D). These results suggest that dietary tryptophan enrichment attenuates LPS-driven inflammatory priming *in vivo*, consistent with suppression of “Signal 1” responses in the lung.

**Figure 4.**
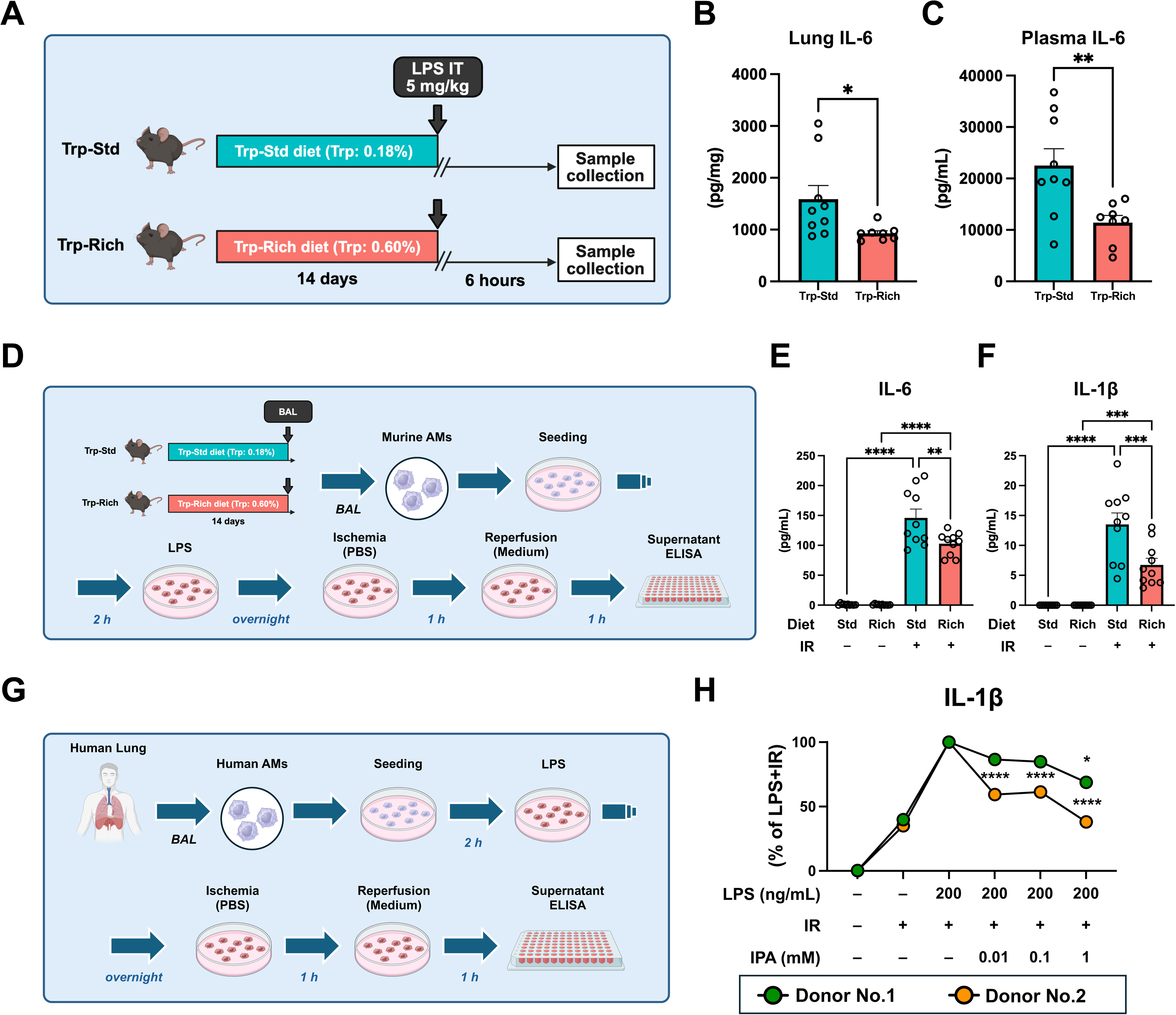
A tryptophan-rich diet suppresses lipopolysaccharide-induced lung inflammation *in vivo*, and reprograms murine and human alveolar macrophage responses *ex vivo*. **A)** Experimental design for the lipopolysaccharide (LPS) intratracheal (IT) challenge. Mice were fed isocaloric tryptophan-standard (Trp-Std, 0.18% tryptophan) or tryptophan-rich (Trp-Rich, 0.60% tryptophan) diets for 2 weeks, followed by LPS IT administration and sample collection 6 h later. **B-C)** A Trp-Rich diet significantly reduced IL-6 levels in systemic plasma and lung tissue after LPS challenge. **D)** Schematic of the *ex vivo* nutritional ischemia-reperfusion (IR) model using primary murine alveolar macrophages (AMs) from mice fed the Trp-Std or Trp-Rich diet for 2 weeks. **E-F)** A Trp-Rich diet significantly reduced IL-6 and IL-1β production in the supernatant after nutritional IR. **G)** Schematic of the *ex vivo* nutritional ischemia-reperfusion model using primary human AMs. **H)** Indole-3-propionate (IPA) reduced IL-1β production in primary human AMs in a dose-dependent manner in two donor lungs. Data in B-C were analyzed by the Mann-Whitney U test (Trp-Std, n = 9; Trp-Rich, n = 8). Data in E-F were analyzed by one-way ANOVA with Tukey’s post hoc test (Trp-Std, n =10; Trp-Rich, n = 10). H were analyzed by one-way ANOVA with Tukey’s post hoc test. *: p<0.05, **: p<0.01, ***: p<0.001, ****: p<0.0001 in (B-C and E-F), *: p<0.05 and ****: p<0.0001 vs LPS+IR in (H).

### A Trp-Rich diet reprograms murine AM responses in an *ex vivo* nutritional IR model

Because our previous work identified AMs as key regulators of lung IR injury, we next focused on AM responses in an *ex vivo* setting [8]. To determine whether dietary tryptophan enrichment reprograms AMs *in vivo*, we isolated primary murine AMs from Trp-Std and Trp-Rich mice after 2 weeks of dietary intervention and subjected them to an *ex vivo* nutritional IR model designed to mimic IR-associated stress by transiently limiting nutrient and serum availability while maintaining normoxic culture conditions (Figure 4D) [21,28]. AMs from Trp-Rich mice released less IL-1β and IL-6 into the supernatant than AMs from Trp-Std mice, indicating that dietary intervention was sufficient to confer a less inflammatory macrophage phenotype *ex vivo* (Figure 4E-F). These findings support the concept that Trp-Rich diet reprograms AM responses *in vivo* and provide an intermediate bridge between the whole-animal phenotype and the downstream *ex vivo* and *in vitro* mechanistic analyses.

### Anti-inflammatory effect of IPA in nutritional IR model with human AMs

Human primary AMs from Donors 1 and 2 were subjected to a nutritional IR model (Figure 4G) [21,28]. Based on characteristic autofluorescence, >90% of the recovered adherent cells were confirmed to be AMs (Figure S6A). Under these conditions, IPA suppressed IL-1β production across 0.01, 0.1, and 1 mM concentrations, and the overall observed effect was consistent across both donors (Figures 4H and S6B-C).

### Comparison of the anti-inflammatory effects of tryptophan metabolites in LPS-stimulated AMs

Given evidence that IPA-AhR signaling can restrain LPS/TLR4/NF-κB “Signal 1” priming, we next used an LPS-driven inflammation model in AMs [8]. Several tryptophan-related metabolites have been reported to exert anti-inflammatory effects, including L-tryptophan, IAA, ILA, indole, and IPA [42–46]. We therefore compared these five metabolites side-by-side. In MH-S cells, pretreatment with each metabolite followed by overnight co-treatment with LPS was performed (Figure 5A). At 0.01 mM, ILA, indole, and IPA significantly reduced IL-6 after LPS stimulation (Figure 5B). At 0.1 mM, all five metabolites reduced IL-6 (Figure 5C). Across both concentrations, IPA produced the largest reduction in IL-6 among the metabolites tested. We then evaluated the same panel in primary human AMs (Donor 3) using overnight co-treatment with LPS and each metabolite (Figure 5D). At 0.01 mM, indole and IPA significantly suppressed IL-6 (Figure 5E). At 0.1 mM, ILA, indole, and IPA reduced IL-6 (Figure 4H). Together, these results identify IPA as the most consistently potent anti-inflammatory metabolite among major tryptophan-related indole derivatives in both murine and human AMs.

**Figure 5.**
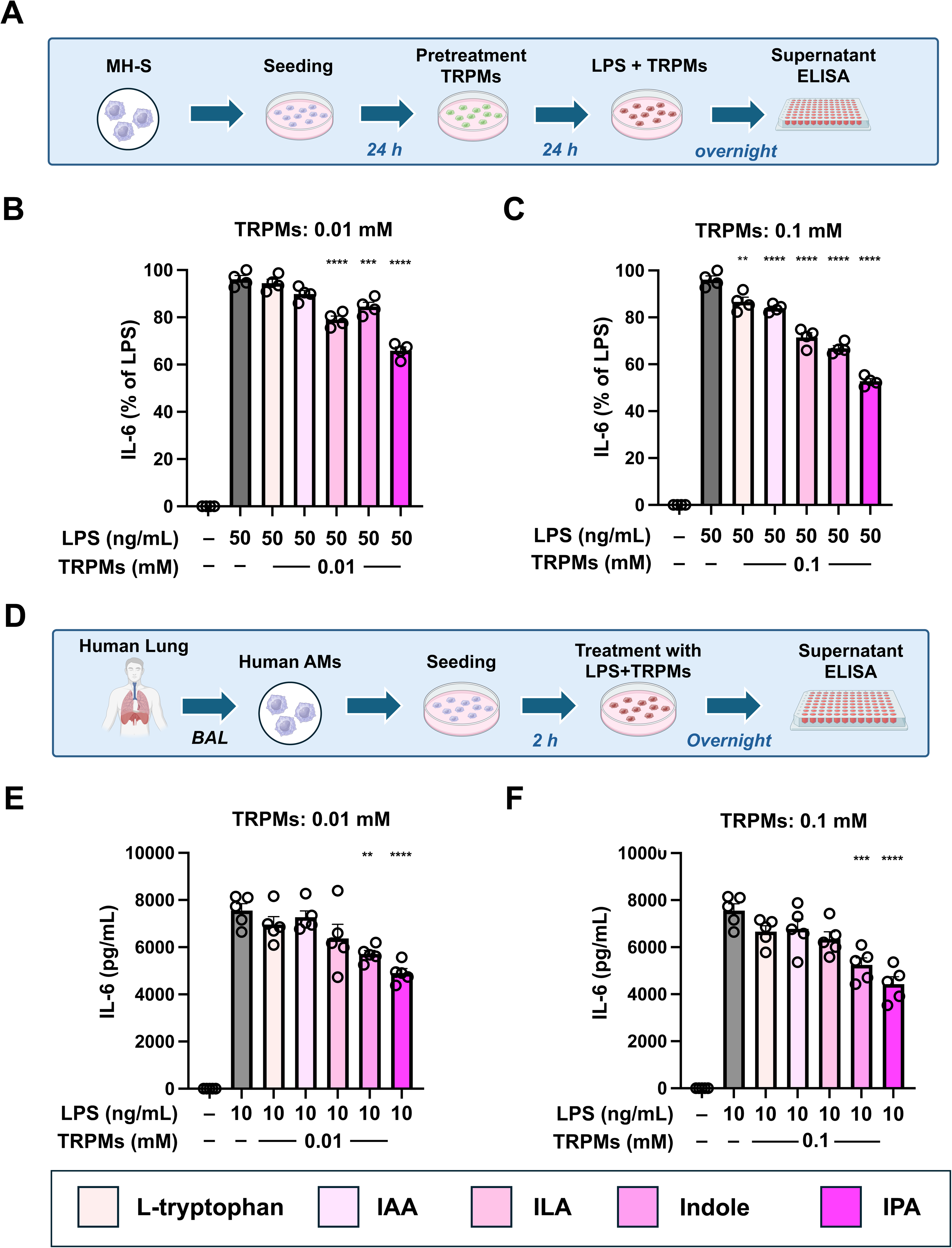
Indole-3-propionate shows the strongest anti-inflammatory effect among representative tryptophan metabolites in murine and human alveolar macrophages. **A)** Experimental design for comparing the anti-inflammatory effects of representative indole metabolites, including L-tryptophan, indole-3-acetate (IAA), indole-3-lactate (ILA), indole, and indole-3-propionate (IPA), in MH-S cells stimulated with lipopolysaccharide (LPS). **B-C)** At 0.01 and 0.1 mM, IPA produced the greatest suppression of IL-6 among the metabolites tested in LPS-stimulated MH-S cells. **D)** Experimental design for comparing representative indole metabolites in primary human alveolar macrophages (AMs) stimulated with LPS. **E-F)** IPA showed the strongest suppression of IL-6 in LPS-stimulated primary human AMs at 0.01 and 0.1 mM. Data were analyzed by one-way ANOVA with Tukey’s post hoc test. **: p<0.01, ***: p<0.001, and ****: p<0.0001 vs LPS alone.

### Anti-inflammatory effect of IPA in AMs

To define the dose-response relationship for IPA, MH-S cells were treated with IPA (0.01, 0.1, and 1 mM) using pretreatment followed by LPS challenge in the continued presence of IPA (Figure 6A). IPA suppressed LPS-induced inflammation in a concentration-dependent manner, whereas IPA alone did not trigger inflammatory activation (Figure 6B). Similar dose-dependent anti-inflammatory effects were observed for L-tryptophan, IAA, ILA, and indole (Figure S7). We next tested IPA in primary human AMs from five donors (Donors 2 and 4-7) using overnight co-treatment with LPS and IPA (Figure 6C). IPA dose-dependently reduced IL-6, IL-1β, CXCL-1, and CXCL-2 in culture supernatants, consistently across donors (Figures 6D-G). In Donor 7, measurable baseline cytokine production was observed in the absence of LPS, consistent with pre-existing inflammatory activation in the donor lung (Figure S8). Notably, IPA reduced IL-1β and CXCL-1 to below baseline in this donor (Figures S8B-C), supporting the possibility that IPA may retain anti-inflammatory activity even in pre-activated AMs. To explore whether baseline AhR pathway tone influenced donor responsiveness to IPA, we related baseline *AHRR* expression to the magnitude of IPA-mediated IL-6 suppression in primary human AMs. AHRR expression was especially higher in Donor No.4 (Figure S9A). In exploratory donor-level analyses, higher baseline *AHRR* expression was associated with greater IPA-mediated reduction of IL-6 at both 0.01 mM and 1 mM IPA (Figures S9B-C). These findings are consistent with the possibility that donor-to-donor variation in baseline AhR pathway tone modulates the magnitude of anti-inflammatory responses to IPA. Collectively, these data show that IPA broadly suppresses cytokine and chemokine production in murine and primary human AMs.

**Figure 6.**
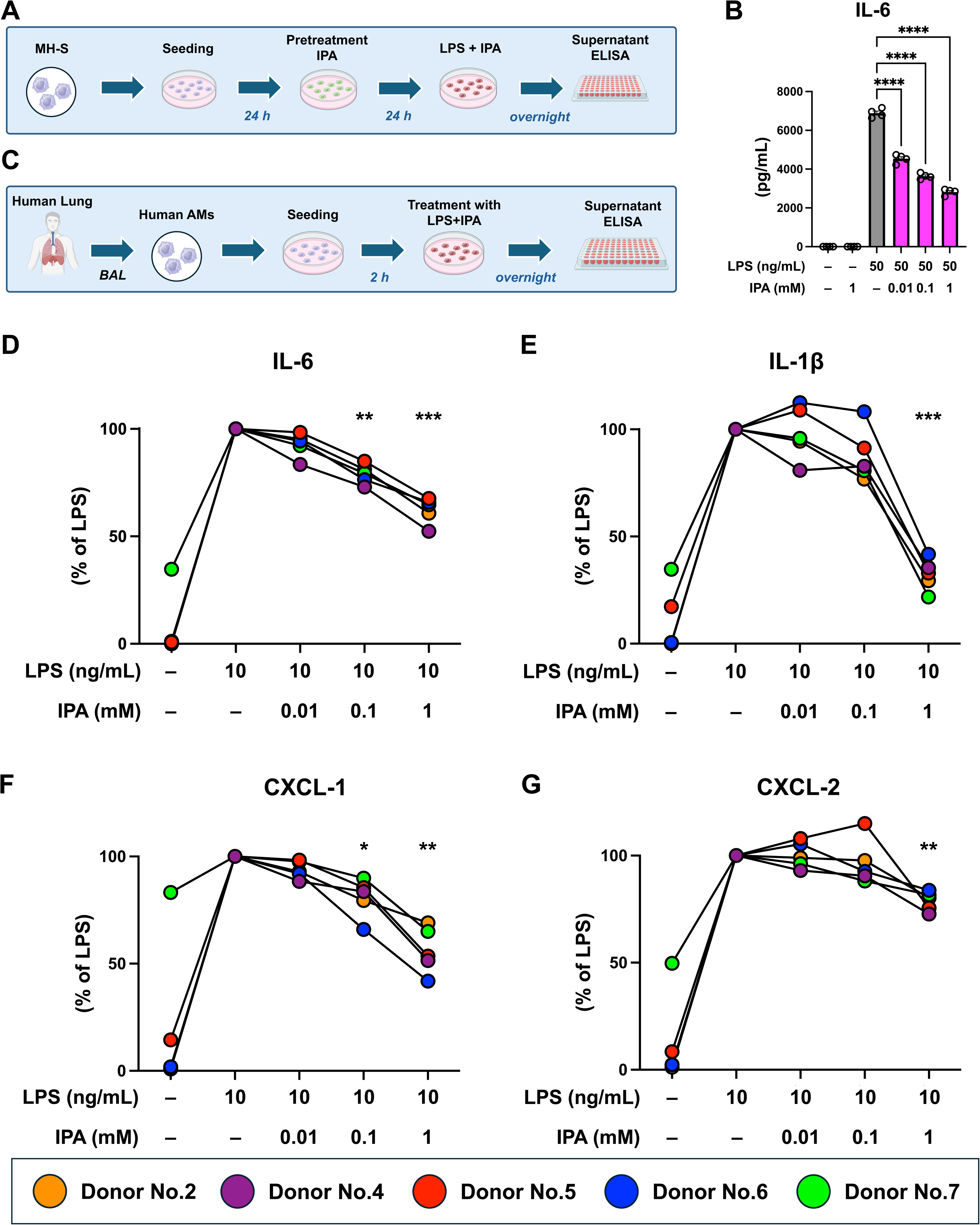
Indole-3-propionate suppresses inflammatory mediator production in murine and human alveolar macrophages in a dose-dependent manner. **A)** Experimental design for evaluating the anti-inflammatory effect of indole-3-propionate (IPA) in MH-S cells stimulated with lipopolysaccharide (LPS). **B)** IPA reduced LPS-induced IL-6 production in a concentration-dependent manner. **C)** Experimental design for evaluating IPA in primary human alveolar macrophages (AMs) stimulated with LPS across five donor lungs. **D-G)** IPA dose-dependently attenuated the production of IL-6, IL-1β, CXCL-1, and CXCL-2 in primary human AMs. Baseline cytokine production was detectable in donor No.7. Notably, at 1 mM IPA, IL-1β and CXCL-1 were reduced below baseline in this donor. Detailed data for donor No.7 are shown in Supplemental Figure S8. Statistical analyses were performed using one-way ANOVA with Tukey’s post hoc test in (B) and repeated-measures one-way ANOVA with Dunnett’s post hoc test (C). *: p<0.05, **: p<0.01, and ***: p<0.001.

### Mechanism of the anti-inflammatory effect of IPA

Because AhR is a major host receptor for microbiota-derived tryptophan metabolites, we tested whether IPA’s anti-inflammatory effects in AMs required AhR signaling. MH-S cells were treated with the AhR antagonist CH-223191, and TCDD was included as a positive-control AhR agonist (Figure 7A). IPA (0.1 mM) and TCDD (10 nM) each suppressed LPS-induced IL-6 production, and these effects were abrogated by CH-223191 (Figure 7B). AhR blockade similarly attenuated IPA-mediated suppression at 0.01 mM, as reflected by IL-6 and TNF-α (Figures S10A-B). We also examined *Ppar*γ, a transcription factor linked to anti-inflammatory programs and homeostatic differentiation in tissue-resident AMs [47,48]. LPS reduced *Ppar*γ expression, whereas IPA restored *Ppar*γ toward baseline (Figure S10C). Together, these findings support an AhR-dependent mechanism for IPA-mediated suppression of inflammatory signaling in AMs and suggest concomitant transcriptional shifts consistent with a more immunoregulatory macrophage state.

**Figure 7.**
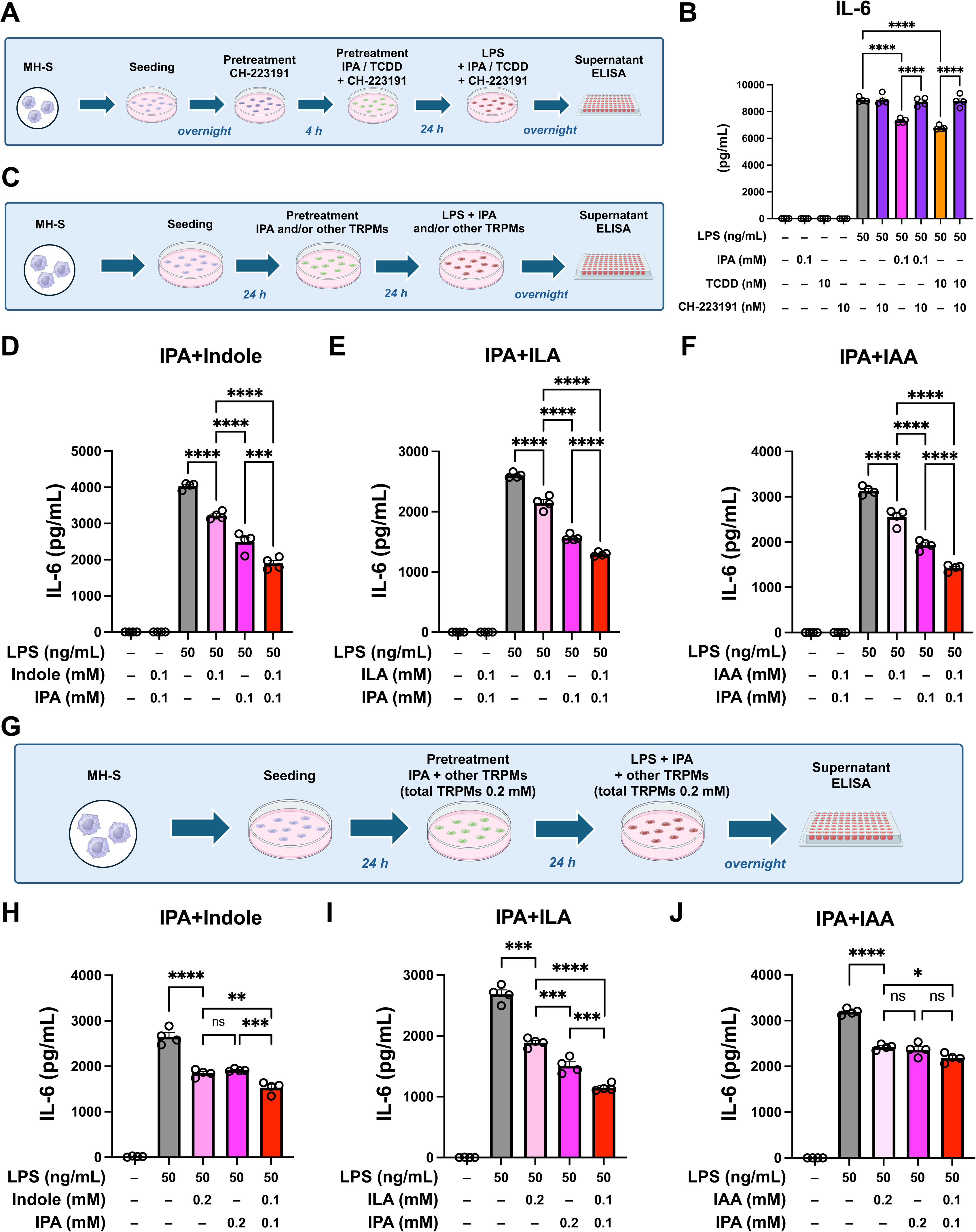
Indole-3-propionate suppresses inflammation via aryl hydrocarbon receptor signaling and exhibits additive and synergistic effects with other indole metabolites. **A)** Experimental design to investigate whether hydrocarbon receptor (AhR) signaling mediates the anti-inflammatory effect of indole-3-propionate (IPA) in MH-S cells stimulated with lipopolysaccharide (LPS). **B)** AhR antagonism with CH-223191 (10 nM) during pre- and co-treatment abolished the suppressive effect of IPA (0.1 mM) on LPS-induced IL-6 production. The AhR agonist 2,3,7,8-tetrachlorodibenzo-p-dioxin (TCDD, 10 nM) served as a positive control. Data for IPA at 0.01 mM are shown in Supplemental Figure S10A-B. **C)** Experimental design to evaluate the additive effect of IPA with other indole metabolites in LPS-stimulated MH-S cells. **D-F)** Co-treatment with IPA and indole, indole-3-lactate (ILA), and indole-3-acetate (IAA) produced greater suppression of LPS-induced IL-6 than either metabolite alone at 0.1 mM. Data at 0.01 and 1 mM are provided in Supplemental Figure S10D-I. **G)** Experimental design to investigate synergistic effect using a fixed total metabolite concentration in LPS-stimulated MH-S cells. **H-J)** At 0.2 mM fixed total concentration, combinations of IPA with indole or ILA suppressed LPS-induced IL-6 more than the corresponding single-metabolite conditions, consistent with synergistic interactions, whereas the IPA+IAA combination did not show a statistically clear synergistic effect. Statistical analysis was performed using one-way ANOVA with Tukey’s post hoc test. *: p<0.05, **: p<0.01, ***: p<0.001, ****: p<0.0001.

### Combined effects of IPA with other tryptophan metabolites

Because IPA naturally coexists with other indole derivatives *in vivo*, we examined whether combining IPA with other tryptophan-related metabolites enhances anti-inflammatory efficacy. In MH-S cells, co-treatment with IPA plus indole, ILA, or IAA produced greater suppression of LPS-induced IL-6 than either IPA or the paired metabolite alone at 0.1 mM (Figures 7C, schema, and 7D-F). Similar additive suppression was observed at 0.01 and 1 mM (Figure S10D-I). To probe potential synergy under a fixed total metabolite exposure, we held the total concentration constant at 0.2 mM, comparing a single metabolite at 0.2 mM with a combination of IPA (0.1 mM) plus a second metabolite (0.1 mM) (Figure 7G). Under these fixed total-concentration conditions, combinations of IPA with indole or ILA suppressed IL-6 more strongly than the corresponding single-metabolite conditions (Figures 7H-I), consistent with synergistic interactions, whereas the IPA+IAA combination did not show a statistically clear synergistic effect (Figure 7J). Overall, these results indicate that IPA can act additively with other tryptophan-derived indoles to dampen AM inflammatory responses.

## Discussion

We investigated whether tryptophan enrichment protects against sterile lung IR injury through the gut–lung axis. A two-week dietary intervention reduced lung inflammation after IR injury and improved oxygenation. *Cyp1a1* and *Cyp1b1* expression increased in lung tissue, suggesting enhanced pulmonary AhR signaling. Targeted metabolomics in feces, PV plasma, and lung tissue revealed a coordinated shift in tryptophan metabolism toward the indole branch, with consistent increases in IPA and decreases in kynurenine/IPA ratios across matrices. In mechanistic studies, IPA showed the strongest anti-inflammatory activity among key tryptophan-derived indoles in MH-S cells and primary human AMs, suppressing multiple cytokines and chemokines in a dose-dependent manner as well as in an *ex vivo* nutritional IR model. AhR blockade reversed IPA-mediated suppression of inflammation, supporting AhR-dependent modulation of macrophage inflammatory programs. Together, these findings are consistent with our model that diet-driven increases in IPA can dampen macrophage “Signal 1” priming upstream of sterile IR injury. Supporting this model *in vivo*, a Trp-Rich diet also reduced LPS-induced IL-6 responses in both lung tissue and systemic plasma.

A Trp-Rich diet reshaped both the gut microbiota composition and the tryptophan metabolite landscape, leading to a consistent increase in indole pathway output in feces, PV plasma, and lung tissue. This rise in IPA is significant because microbiota-derived tryptophan metabolites can engage host AhR-dependent programs that support barrier homeostasis and immunoregulatory functions, providing a mechanistic rationale for gut-to-distal organ communication [49]. Notably, *Lactobacillus* and *Bifidobacterium* expanded in the Trp-Rich group, and these taxa are linked to indole-related metabolism through production of downstream indole derivatives, which can shape host immune tone [50,51]. Accordingly, the integrated analyses linked higher IPA levels to lung inflammatory readouts that reflect reduced inflammatory activity and engagement of the AhR pathway. The biological plausibility of focusing on IPA was further supported by *in vivo* studies from other groups showing that IPA engages host immunoregulatory programs, including AhR-dependent macrophage functions, and can attenuate inflammatory injury, aligning with our observation that IPA was the most potent anti-inflammatory indole among those tested on AMs in the context of IR and LPS sterile injury [23,52,53].

Building on our observations of multi-compartmental increases in IPA along the gut-lung axis, we next investigated whether IPA could directly reprogram AM inflammatory responses in a manner consistent with our *in vivo* dietary intervention observations. Consistent with reduced inflammatory priming *in vivo*, a complementary *in vivo* LPS intratracheal challenge further supported this phenotype and provided a translational bridge to our AM-focused mechanistic assays. Across both MH-S cells and primary human AMs, IPA showed the most potent suppression of LPS-induced cytokine and chemokine production among the indole derivatives tested, with clear dose dependence across physiologic-to-pharmacologic concentration ranges. These results align with prior work showing that AhR signaling can restrain TLR4-driven cytokine responses in macrophages, and that microbiota-derived IPA can act as an AhR ligand, shaping macrophage effector functions and inflammatory injury [54]. Mechanistically, pharmacologic AhR blockade with CH-223191 abrogated the anti-inflammatory effect of IPA, while the AhR agonist TCDD recapitulated suppression of IL-6, supporting an AhR-dependent mode of action [23]. Moreover, IPA restored *Ppar*γ expression in LPS-stimulated MH-S cells, consistent with established roles for PPARγ in AM differentiation and homeostatic programs [48]. Since IPA coexists with other indole derivatives *in vivo*, our observation that combinations of IPA with indole or ILA further enhanced suppression of IL-6 is conceptually consistent with the literature showing that distinct microbiota-derived indoles can converge on host immunoregulatory pathways [55,56]. From a translational perspective, our data support the concept that perioperative “dietary prehabilitation” targeting microbial tryptophan metabolism could be a feasible way to protect the lung against sterile inflammatory injury, given that IR risk is often predictable in transplantation and cardiothoracic surgery. Consistent with the airway relevance of this metabolic axis, reduced microbiota-derived IPA has been linked to worse allergic airway inflammation *in vivo*, supporting the idea that IPA availability can shape lung immune tone [46]. Additionally, tryptophan supplementation attenuated LPS-induced acute lung injury in rats, providing additional rationale for leveraging dietary tryptophan availability to modulate lung inflammation [42]. Beyond diet alone, a more controllable strategy may be microbiome-directed therapy that directly elevates endogenous IPA. In clinical contexts with predictable inflammatory risk, disruption of gut microbial IPA production has been associated with worse postoperative outcomes [57]. Restoring IPA signaling conferred protective effects, whereas colonization with an IPA-deficient *Clostridium sporogenes* strain abrogated protection [57]. Complementing this, *Clostridium sporogenes*-derived metabolites (including IPA) have been shown to protect against colonic inflammation, supporting the feasibility of leveraging defined IPA-producing strains as next-generation probiotics [53]. Synbiotic design may further optimize this approach, as pairing *Clostridium sporogenes* with a matching dietary substrate increased gut-derived IPA levels and improved disease phenotypes *in vivo* [58]. More broadly, dietary fiber composition can redirect community tryptophan metabolism toward higher IPA production via microbial interactions, suggesting that diet composition and strain selection should be co-designed rather than considered separately [59]. Finally, this gut–lung axis model offers practical pharmacodynamic readouts, such as circulating IPA and the kynurenine/IPA ratio, to assess whether interventions engage the intended gut-to-lung metabolic axis in humans.

Several limitations should be considered. First, our *in vivo* study was powered to detect diet-associated differences but used a modest sample size and ultimately focused on male mice because the protective phenotype was not observed in females. Sex hormones can modulate lung IR biology, and future sex-stratified studies will be necessary [34]. Second, while targeted metabolomics showed reductions in the kynurenine/IPA and kynurenine/tryptophan ratios, these ratios should be interpreted as indirect surrogates of pathway balance and immune-linked tryptophan catabolism rather than direct measures of metabolic flux. Complementary approaches that quantify kinetics or pathway activity in relevant compartments will strengthen mechanistic interpretation [60,61]. Third, our *in vitro* and *ex vivo* experiments intentionally spanned putative physiologic-to-pharmacologic concentrations of IPA, but the practical exposure of alveolar lining fluid and tissue-resident AMs to tryptophan metabolites *in vivo* remains uncertain, motivating future pharmacokinetic and dosing studies. Finally, although pharmacologic AhR antagonism supported an AhR-dependent mechanism, cell-specific genetic approaches (e.g., myeloid- or AM-targeted AhR perturbation) and *in vivo* intervention studies will be needed to establish causality across the gut–lung axis [62].

### Conclusions

A Trp-Rich diet attenuated sterile lung IR injury, coinciding with gut microbiota remodeling, a systemic and pulmonary shift in tryptophan metabolism toward indole production, and increased endogenous IPA levels across feces, PV plasma, and lung tissue. Consistent with lower lung immune tone, the Trp-Rich diet also dampened LPS-induced IL-6 responses *in vivo*. IPA showed robust anti-inflammatory activity in murine and human AMs, acted through AhR-dependent signaling, and was enhanced by combining it with other indole metabolites. Together, these findings support dietary modulation of microbial tryptophan metabolism as a feasible strategy to raise lung IPA levels and reprogram macrophage responses to suppress acute lung IR inflammation.

## Acknowledgements

Metabolomics measurements were performed by the University of Michigan Medical School BRCF Metabolomics Core Facility under the supervision of Maureen Kachmann (RRID: SCR_026721). The human lungs were generously provided for research purposes by Dr. Michael Matthay in the Department of Medicine and Anesthesia at the University of California San Francisco. 16S rRNA sequencing was performed by Dr. Rahim Khan in the Benioff Center for Microbiome Medicine (BCMM) at the University of California San Francisco.

## Author contributions

TC designed the study, conducted the study, collected the data, analyzed the data, wrote the manuscript, and revised the manuscript. DM, TD and TX contributed to the study design, conducted experiments, analyzed the data, and revised the manuscript. AP supervised the study, contributed to study design, interpreted all data, and revised the manuscript.

## Ethics approval and informed consent

All animal studies were approved by the Institutional Animal Care and Use Committee at the University of California San Francisco (IACUC protocol: AN197325-01). Explicit approval for the use of donor lungs for research from each donor’s family was obtained by Donor Network West, and the UCSF Biosafety Committee approved the experiments using cadaver human lung tissue.

## Consent for publication

Not applicable.

## Availability of data and materials

The datasets generated during and/or analyzed during the current study are available from the corresponding author on reasonable request.

