## Supplemental Figure Legends for "Dietary tryptophan mitigates lung ischemia-reperfusion injury via microbiota-derived indole-3-propionate and aryl hydrocarbon receptor signaling"

**Supplemental Figure Legend**

**Figure S1. Effect of a tryptophan-rich diet on body weight and sex-stratified lung inflammation. A)** Body weight changes during the 2-week dietary intervention with a tryptophan-standard (Trp-Std) and tryptophan-rich (Trp-Rich) diet. **B)** Representative gross appearance of the intestinal tract in the Trp-Std and Trp-Rich groups. **C)** Lung IL-1β and **D)** IL-6 after lung ischemia-reperfusion injury, stratified by sex, showing that the anti-inflammatory effect of a Trp-Rich diet was evident in male mice but not in female mice. Statistical comparisons were performed using an unpaired t-test. Male Trp-Std, n = 12, female Trp-Std, n = 11; male Trp-Rich, n = 6; female Trp-Rich, n = 9. *: p<0.05.

**Figure S2. Remodeling of gut microbiota by a Trp-Rich diet.** **A-D)** Additional fecal microbiota alpha-diversity metrics (Chao1, ACE, Simpson, and Fisher indices), indicating reduced diversity in the tryptophan-rich (Trp-Rich) group. **E)** *Genus*-level taxonomic composition. The top 12 *genera* are shown. I) Species-level differential abundance visualized by a volcano plot. *Lactobacillus intestinalis* was significantly and markedly increased in the Trp-Rich group. Statistical comparisons were performed using the Mann-Whitney test with false discovery rate correction for multiple testing. n = 4 in each group.

**Figure S3. Comprehensive targeted tryptophan metabolite profiling in feces, PV plasma, and portal vein plasma. A)** Box plots displaying the distribution of individual feces tryptophan metabolites in each group. Kynurenic acid and 5-hydroxyindoleacetic acid were significantly increased in the tryptophan-rich (Trp-Rich) group. **B)** Box plots displaying the distribution of individual portal vein plasma tryptophan metabolites in each group. **C)** Box plots displaying the distribution of individual lung tryptophan metabolites in each group. Kynurenine was significantly increased in the Trp-Rich group. Statistical comparisons were performed using the Mann-Whitney test with false discovery rate correction for multiple testing. Trp-Std, n = 10; Trp-Rich, n = 6.

**Figure S4. Integrated correlation analyses linking gut taxa, indole-3-propionate, and aryl hydrocarbon receptor pathway readouts.** **A)** *Genus*-level correlations between gut microbiota and lung inflammatory markers showing *Bifidobacterium* abundance was inversely correlated with lung IL-6. **B-C)** Correlation analyses between the relative abundance of *Bifidobacterium* or *Lactobacillus* and inflammatory cytokines (IL-1β, IL-6) or aryl hydrocarbon receptor (AhR) pathway readouts (*Cyp1a1*, *Cyp1b1*). **D)** *Species*-level correlations showing that *Lactobacillus intestinalis* abundance was inversely correlated with lung IL-6. **E)** Correlation analyses between the relative abundance of *Lactobacillus intestinalis* and IL-1β, IL-6, *Cyp1a1*, or *Cyp1b1*. **F)** Correlation analyses between selected gut microbial taxa (*Bifidobacterium*, *Lactobacillus*, and *Lactobacillus intestinalis*) and indole-3-propionate (IPA) concentrations in feces, portal vein (PV) plasma, and lung tissue. **G)** Association between lung *Cyp1a1* expression and IPA concentration in PV plasma. **H)** Association between Lung *Cyp1a1* and lung IPA concentration. Correlations were assessed using Spearman’s rank correlation. Trp-Std, n = 10; Trp-Rich, n = 6. In microbiome-related analysis, n = 4 in each group.

**Figure S5. Effect of tryptophan-rich diet on lipopolysaccharide-induced lung injury. A)** *Il6* gene expression, **B)** IL-1β level, **C)** *Il1β* gene expression, and **D)** TNF-α level in lung tissue 6 h after intratracheal lipopolysaccharide (LPS) challenge. Data were analyzed using the Mann-Whitney test. *: p<0.05.

**Figure S6. Indole-3-propionate suppresses inflammatory cytokine production in an *ex vivo* nutritional ischemia-reperfusion injury model using primary human alveolar macrophages. A)** Representative image of autofluorescence of primary human alveolar macrophages (AMs). **B-C)** Primary human AMs from donor lungs No.1 and No.2 were evaluated in the *ex vivo* nutritional ischemia-reperfusion model. Indole-3-propionate (IPA) reduced IL-1β production in a concentration-dependent manner. Data were analyzed by one-way ANOVA with Tukey’s post hoc test. *: p<0.05 and ****: p<0.0001 vs LPS+IR.

**Figure S7. Dose-dependent anti-inflammatory effects of representative indole metabolites in lipopolysaccharide-stimulated MH-S cells.** In MH-S cells stimulated with lipopolysaccharide (LPS), **A)** L-tryptophan, **B)** indole-3-acetate, **C)** indole-3-lactate, and **D)** indole each attenuated IL-6 production in a concentration-dependent manner. The 1 mM L-tryptophan condition was not tested due to limited solubility. Statistical analyses were performed by one-way ANOVA with Tukey’s post hoc test. *: p<0.05, **: p<0.01, ***: p<0.001, and ****: p<0.0001.

**Figure S8. Indole-3-propionate suppresses inflammatory mediator production in lipopolysaccharide-stimulated primary human alveolar macrophages from donor No.7.** Primary human alveolar macrophages were stimulated with lipopolysaccharide and treated with Indole-3-propionate (IPA). IPA reduced **A)** IL-6, **B)** IL-1β, **C)** CXCL-1, and **D)** CXCL-2 in a concentration-dependent manner. Baseline cytokine/chemokine production was detectable in this donor. Notably, IL-1β and CXCL-1 were reduced below baseline at 1 mM IPA concentration. Statistical analyses were performed using one-way ANOVA with Tukey’s post hoc test. *: p<0.05, **: p<0.01, ***: p<0.001, and ****: p<0.0001.

**Figure S9. Baseline AHRR expression is associated with donor responsiveness to indole-3-propionate in primary human alveolar macrophages. A)** *AHRR* gene expression in human primary alveolar macrophages (AMs) from donors No.2 and 4-7. *AHRR* expression was extremely high in donor No.4. **B-C)** Correlations between AHRR gene expression in AMs and IL-6 reduction relative to the lipopolysaccharide-alone condition at 0.01 and 1 mM indole-3-propionate. AHRR expression was significantly positively correlated with IL-6 reduction in the supernatant. Data was analyzed by Pearson’s correlation analysis.

**Figure S10. Aryl hydrocarbon receptor dependence of indole-3-propionate-mediated anti-inflammatory effects and additive interactions with other indole metabolites in lipopolysaccharide-stimulated MH-S cells. A-B)** MH-S cells were stimulated with lipopolysaccharide (LPS) and treated with indole-3-propionate (IPA, 0.01 mM) with or without pre-and co-treatment with the AhR antagonist CH-223191 (10 nM). Ch-223191 abolished IPA-mediated suppression of IL-6 and TNF-α. **C)** IPA pretreatment restored the *Pparγ* gene expression that was reduced by LPS stimulation. **D-I)** Additive effects of IPA in combination with indole, indole-3-lactate (ILA), and indole-3-acetate (IAA) at 0.01 and 1 mM concentrations on LPS-induced IL-6 production. Statistical analyses were performed using one-way ANOVA with Tukey’s post hoc test. *: p<0.05, **: p<0.01, ***: p<0.001, and ****: p<0.0001.
