## Supplemental Figures for "Dietary tryptophan mitigates lung ischemia-reperfusion injury via microbiota-derived indole-3-propionate and aryl hydrocarbon receptor signaling"

### Slide 1
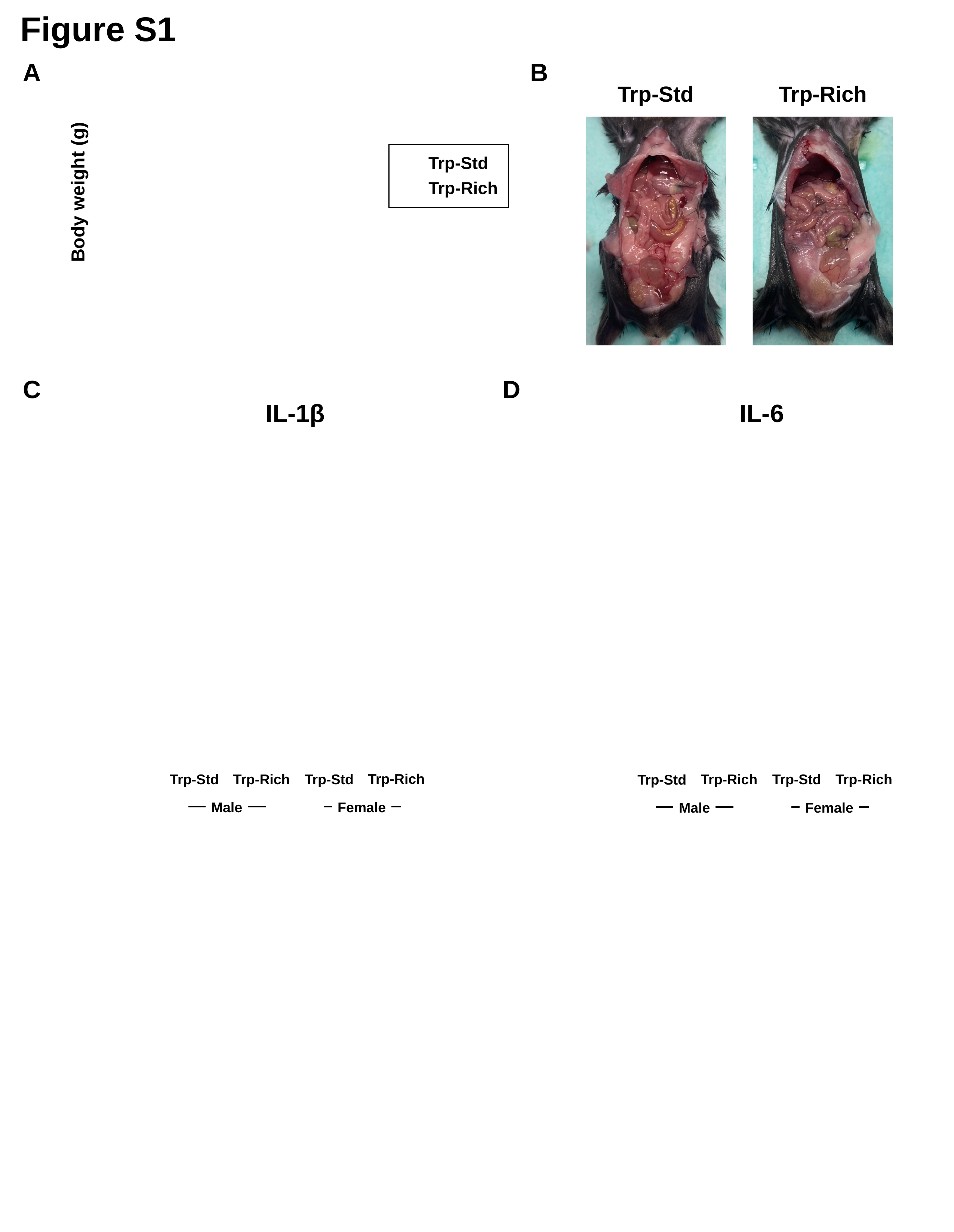

Figure S1
B
A
Trp-Std
Trp-Rich
Trp-Std
Trp-Rich
Body weight (g)
C
D
IL-1β
Trp-Rich
Trp-Rich
Trp-Std
Trp-Std
Male
Female
IL-6
Trp-Rich
Trp-Rich
Trp-Std
Trp-Std
Male
Female

### Slide 2
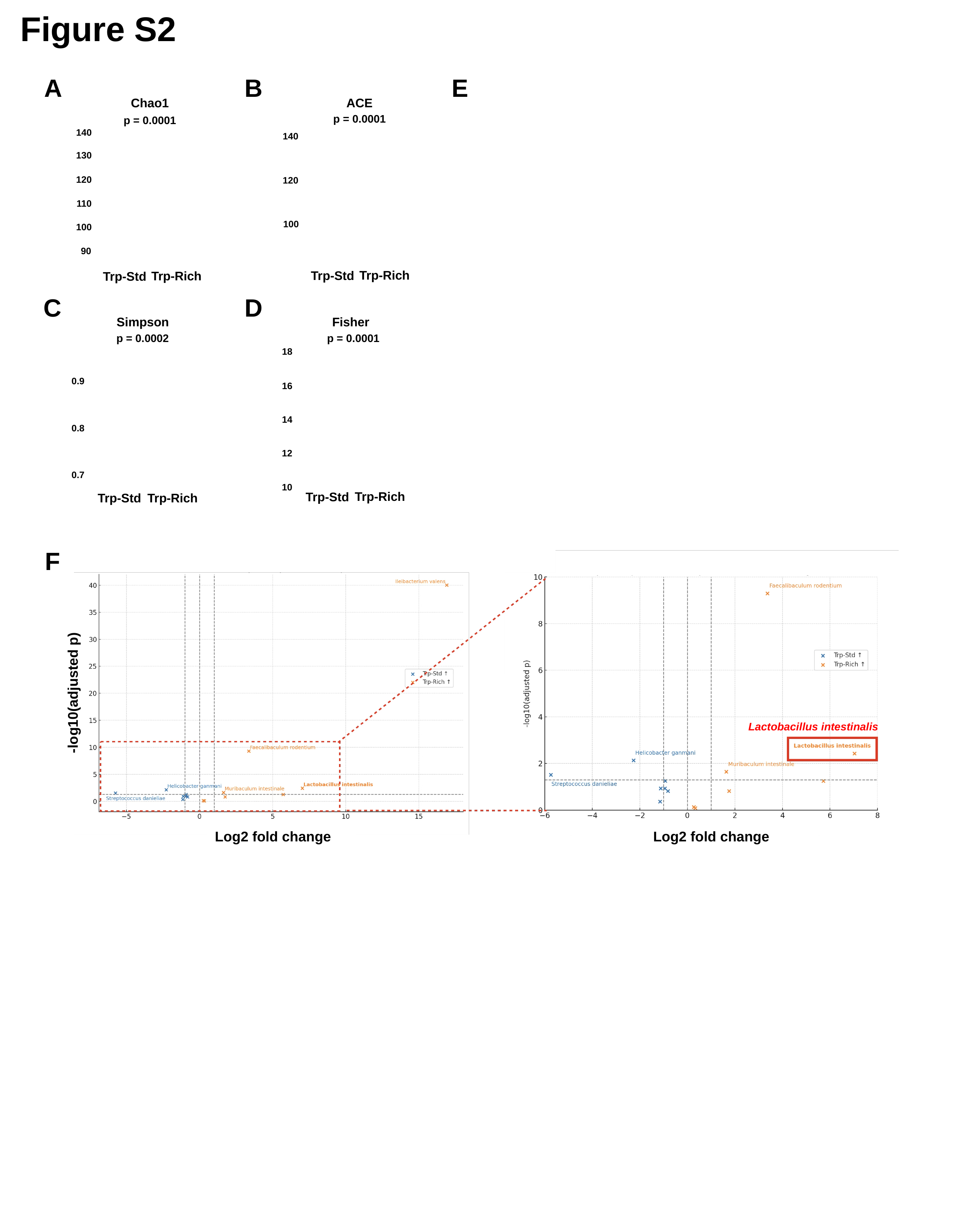

Figure S2
B
E
A
Chao1
p = 0.0001
140
130
120
110
100
90
Trp-Rich
Trp-Std
ACE
p = 0.0001
140
120
100
Trp-Rich
Trp-Std
C
D
Fisher
p = 0.0001
18
16
14
12
10
Trp-Rich
Trp-Std
Simpson
p = 0.0002
0.9
0.8
0.7
Trp-Rich
Trp-Std
F
-log10(adjusted p)
Lactobacillus intestinalis
Log2 fold change
Log2 fold change

### Slide 3
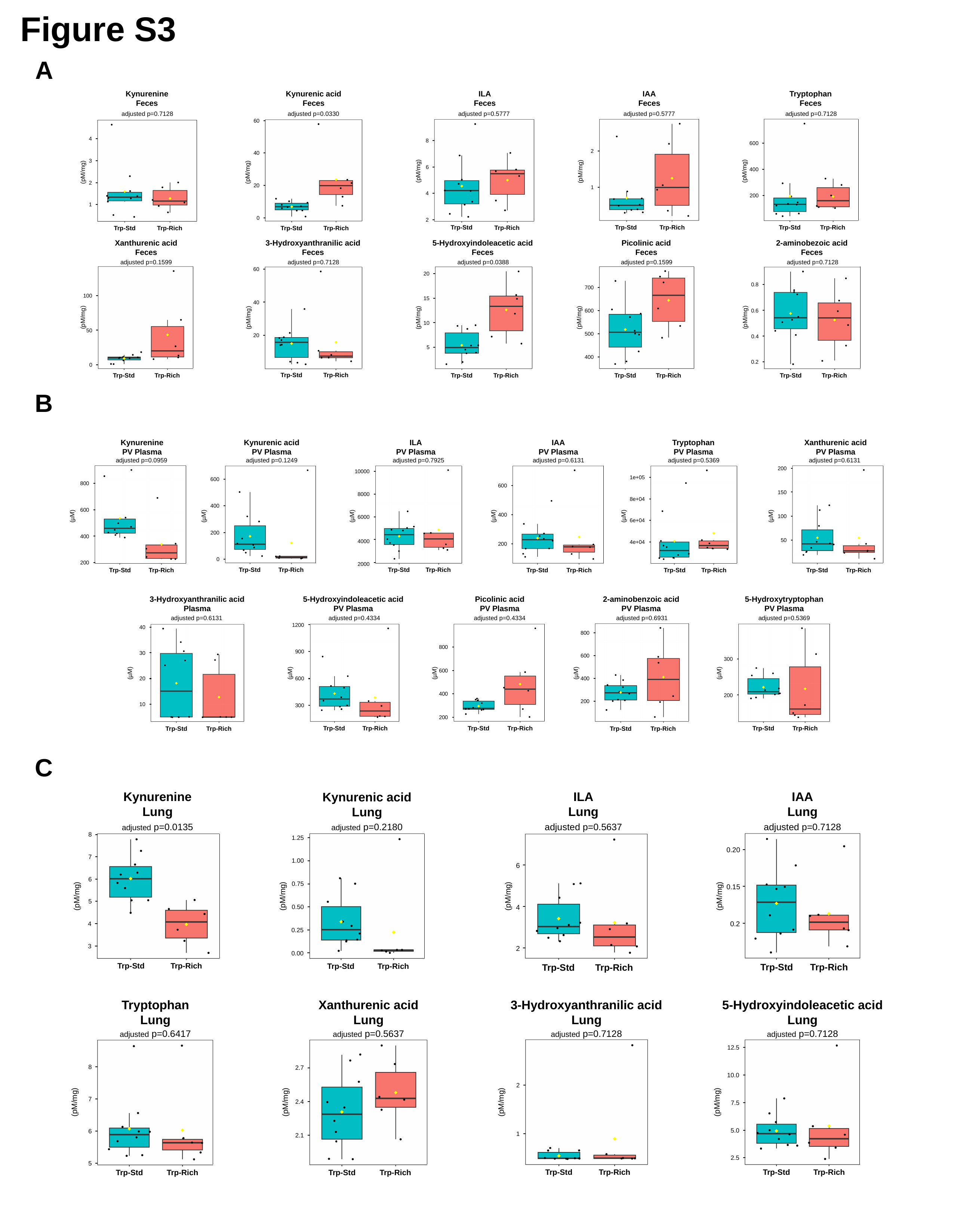

Figure S3
A
Kynurenine
Feces
adjusted p=0.7128
4
3
(pM/mg)
2
1
Trp-Rich
Trp-Std
Kynurenic acid
Feces
adjusted p=0.0330
60
40
(pM/mg)
20
0
Trp-Rich
Trp-Std
ILA
Feces
adjusted p=0.5777
8
6
(pM/mg)
4
2
Trp-Std
Trp-Rich
IAA
Feces
adjusted p=0.5777
2
1
(pM/mg)
Trp-Std
Trp-Rich
Tryptophan
Feces
adjusted p=0.7128
600
400
(pM/mg)
200
Trp-Rich
Trp-Std
Xanthurenic acid
Feces
adjusted p=0.1599
100
(pM/mg)
50
0
Trp-Rich
Trp-Std
3-Hydroxyanthranilic acid
Feces
adjusted p=0.7128
60
40
(pM/mg)
20
Trp-Rich
Trp-Std
5-Hydroxyindoleacetic acid
Feces
adjusted p=0.0388
20
15
(pM/mg)
10
5
Trp-Rich
Trp-Std
Picolinic acid
Feces
adjusted p=0.1599
700
600
(pM/mg)
500
400
Trp-Rich
Trp-Std
2-aminobezoic acid
Feces
adjusted p=0.7128
0.8
0.6
(pM/mg)
0.4
0.2
Trp-Rich
Trp-Std
B
Kynurenine
PV Plasma
adjusted p=0.0959
800
600
(µM)
400
200
Trp-Rich
Trp-Std
Kynurenic acid
PV Plasma
adjusted p=0.1249
600
400
(µM)
200
0
Trp-Rich
Trp-Std
ILA
PV Plasma
adjusted p=0.7925
10000
8000
(µM)
6000
4000
2000
Trp-Rich
Trp-Std
IAA
PV Plasma
adjusted p=0.6131
600
400
(µM)
200
Trp-Rich
Trp-Std
Tryptophan
PV Plasma
adjusted p=0.5369
1e+05
8e+04
(µM)
6e+04
4e+04
Trp-Rich
Trp-Std
Xanthurenic acid
PV Plasma
adjusted p=0.6131
200
150
(µM)
100
50
Trp-Rich
Trp-Std
3-Hydroxyanthranilic acid
Plasma
adjusted p=0.6131
40
30
(µM)
20
10
Trp-Rich
Trp-Std
5-Hydroxyindoleacetic acid
PV Plasma
adjusted p=0.4334
1200
900
(µM)
600
300
Trp-Rich
Trp-Std
Picolinic acid
PV Plasma
adjusted p=0.4334
800
600
(µM)
400
200
Trp-Rich
Trp-Std
2-aminobenzoic acid
PV Plasma
adjusted p=0.6931
800
600
(µM)
400
200
Trp-Rich
Trp-Std
5-Hydroxytryptophan
PV Plasma
adjusted p=0.5369
300
(µM)
200
Trp-Rich
Trp-Std
C
IAA
Lung
adjusted p=0.7128
0.20
0.15
(pM/mg)
0.2
Trp-Std
Trp-Rich
ILA
Lung
adjusted p=0.5637
6
(pM/mg)
4
2
Trp-Rich
Trp-Std
Kynurenine
Lung
adjusted p=0.0135
8
7
6
(pM/mg)
5
4
3
Trp-Rich
Trp-Std
Kynurenic acid
Lung
adjusted p=0.2180
1.25
1.00
0.75
(pM/mg)
0.50
0.25
0.00
Trp-Rich
Trp-Std
Tryptophan
Lung
adjusted p=0.6417
8
7
(pM/mg)
6
5
Trp-Rich
Trp-Std
Xanthurenic acid
Lung
adjusted p=0.5637
2.7
(pM/mg)
2.4
2.1
Trp-Rich
Trp-Std
3-Hydroxyanthranilic acid
Lung
adjusted p=0.7128
2
(pM/mg)
1
Trp-Rich
Trp-Std
5-Hydroxyindoleacetic acid
Lung
adjusted p=0.7128
12.5
10.0
(pM/mg)
7.5
5.0
2.5
Trp-Rich
Trp-Std

### Slide 4
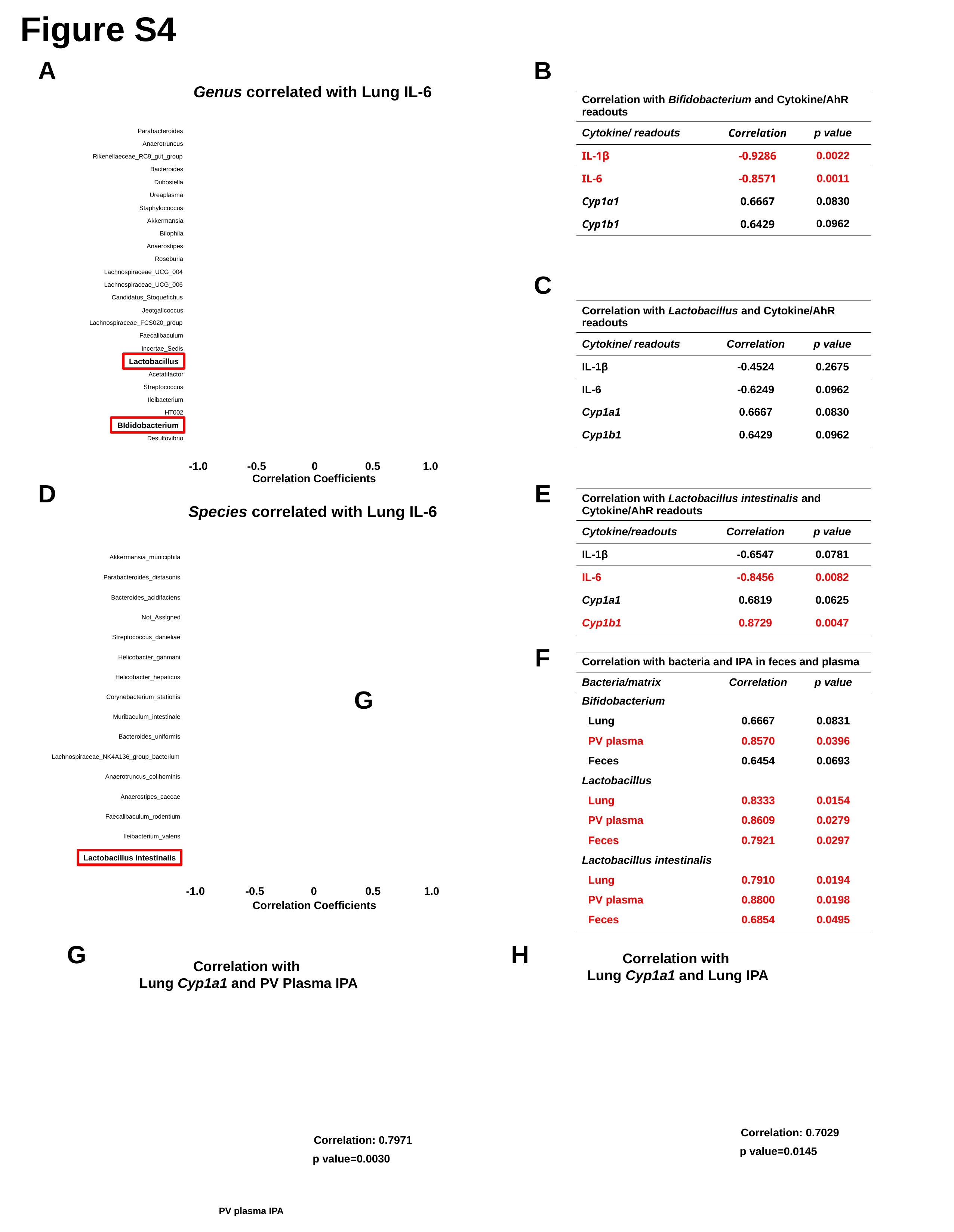

Figure S4
A
B
Genus correlated with Lung IL-6
Correlation Coefficients
Lactobacillus
BIdidobacterium
-1.0
-0.5
0
0.5
1.0
Parabacteroides
Anaerotruncus
Rikenellaeceae_RC9_gut_group
Bacteroides
Dubosiella
Ureaplasma
Staphylococcus
Akkermansia
Bilophila
Anaerostipes
Roseburia
Lachnospiraceae_UCG_004
Lachnospiraceae_UCG_006
Candidatus_Stoquefichus
Jeotgalicoccus
Lachnospiraceae_FCS020_group
Faecalibaculum
Incertae_Sedis
Acetatifactor
Streptococcus
Ileibacterium
HT002
Desulfovibrio
| Correlation with Bifidobacterium and Cytokine/AhR readouts | | |
| --- | --- | --- |
| Cytokine/ readouts | Correlation | p value |
| IL-1β | -0.9286 | 0.0022 |
| IL-6 | -0.8571 | 0.0011 |
| Cyp1a1 | 0.6667 | 0.0830 |
| Cyp1b1 | 0.6429 | 0.0962 |
C
| Correlation with Lactobacillus and Cytokine/AhR readouts | | |
| --- | --- | --- |
| Cytokine/ readouts | Correlation | p value |
| IL-1β | -0.4524 | 0.2675 |
| IL-6 | -0.6249 | 0.0962 |
| Cyp1a1 | 0.6667 | 0.0830 |
| Cyp1b1 | 0.6429 | 0.0962 |
D
E
| Correlation with Lactobacillus intestinalis and Cytokine/AhR readouts | | |
| --- | --- | --- |
| Cytokine/readouts | Correlation | p value |
| IL-1β | -0.6547 | 0.0781 |
| IL-6 | -0.8456 | 0.0082 |
| Cyp1a1 | 0.6819 | 0.0625 |
| Cyp1b1 | 0.8729 | 0.0047 |
Species correlated with Lung IL-6
Lactobacillus intestinalis
-1.0
-0.5
0
0.5
1.0
Correlation Coefficients
Akkermansia_municiphila
Parabacteroides_distasonis
Bacteroides_acidifaciens
Not_Assigned
Streptococcus_danieliae
Helicobacter_ganmani
Helicobacter_hepaticus
Corynebacterium_stationis
Muribaculum_intestinale
Bacteroides_uniformis
Lachnospiraceae_NK4A136_group_bacterium
Anaerotruncus_colihominis
Anaerostipes_caccae
Faecalibaculum_rodentium
Ileibacterium_valens
F
| Correlation with bacteria and IPA in feces and plasma | | |
| --- | --- | --- |
| Bacteria/matrix | Correlation | p value |
| Bifidobacterium | | |
| Lung | 0.6667 | 0.0831 |
| PV plasma | 0.8570 | 0.0396 |
| Feces | 0.6454 | 0.0693 |
| Lactobacillus | | |
| Lung | 0.8333 | 0.0154 |
| PV plasma | 0.8609 | 0.0279 |
| Feces | 0.7921 | 0.0297 |
| Lactobacillus intestinalis | | |
| Lung | 0.7910 | 0.0194 |
| PV plasma | 0.8800 | 0.0198 |
| Feces | 0.6854 | 0.0495 |
G
G
H
Correlation with
Lung Cyp1a1 and Lung IPA
Correlation: 0.7029
p value=0.0145
Correlation with
Lung Cyp1a1 and PV Plasma IPA
Correlation: 0.7971
p value=0.0030
PV plasma IPA

### Slide 5
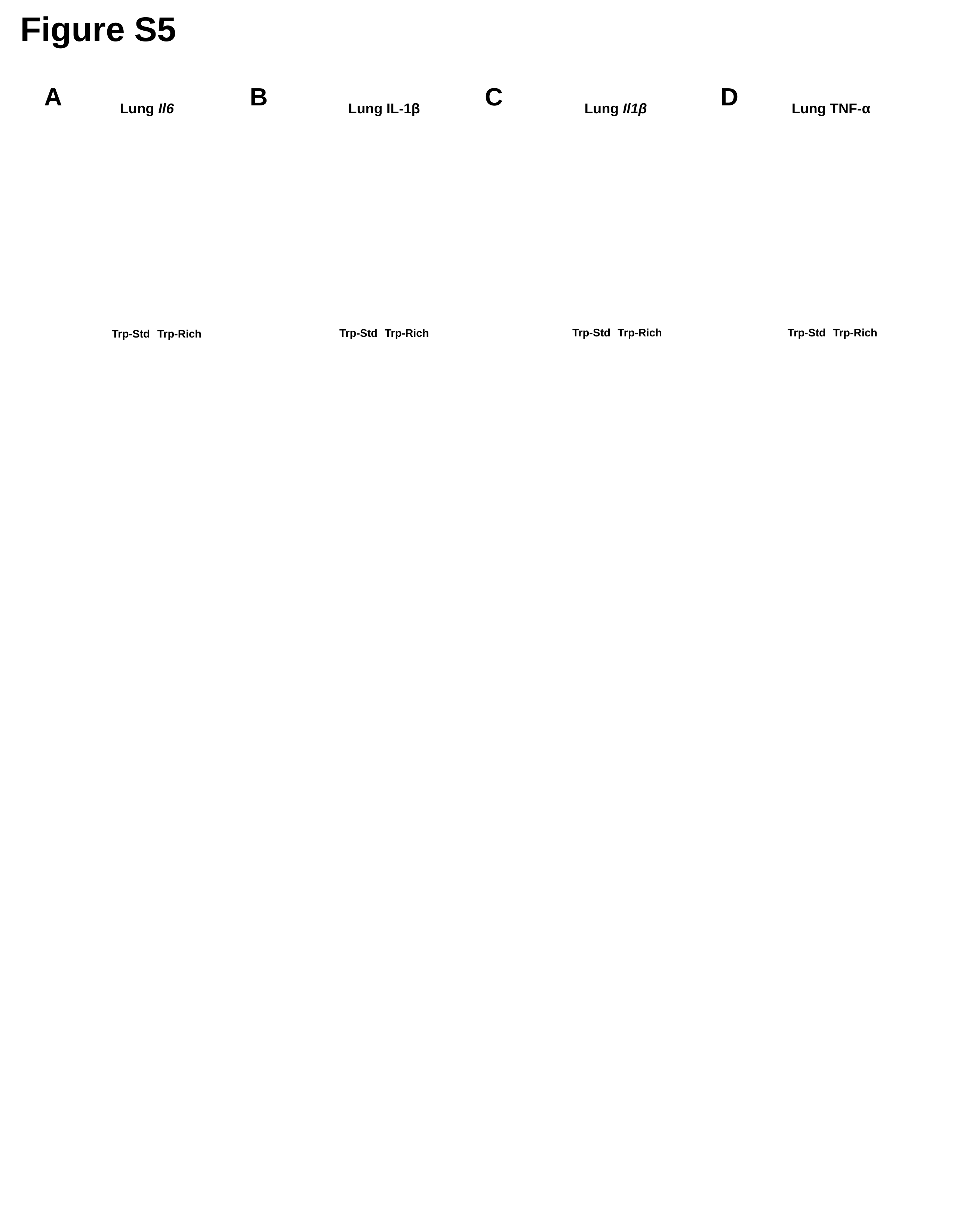

Figure S5
A
B
C
D
Lung Il6
Trp-Rich
Trp-Std
Lung IL-1β
Trp-Rich
Trp-Std
Lung Il1β
Trp-Std
Trp-Rich
Lung TNF-α
Trp-Std
Trp-Rich

### Slide 6
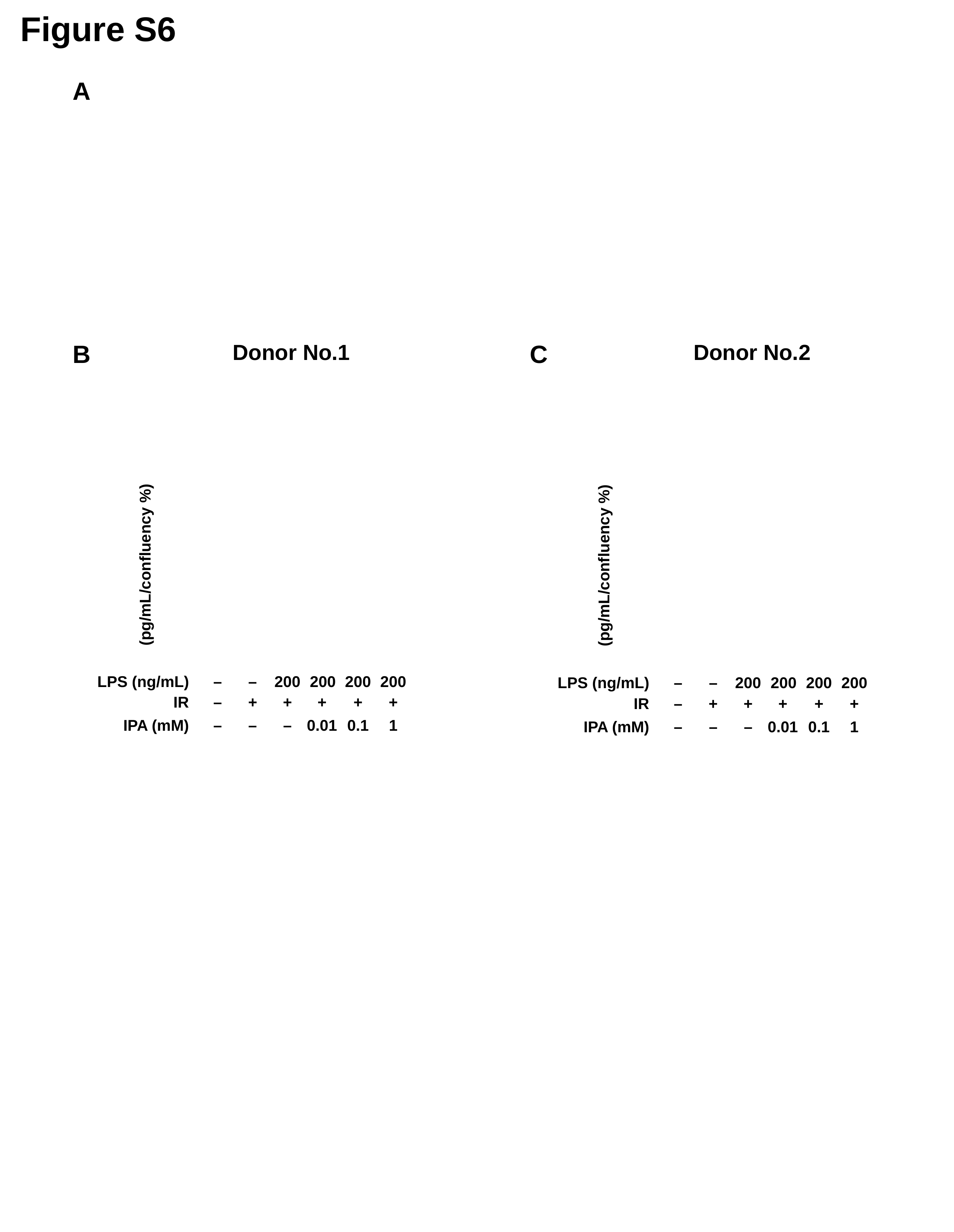

Figure S6
A
Donor No.1
(pg/mL/confluency %)
–
–
200
200
200
200
+
+
IR
–
+
+
+
1
0.1
IPA (mM)
–
–
–
0.01
LPS (ng/mL)
B
Donor No.2
(pg/mL/confluency %)
LPS (ng/mL)
–
–
200
200
200
200
+
+
IR
–
+
+
+
1
0.1
IPA (mM)
–
–
–
0.01
C

### Slide 7
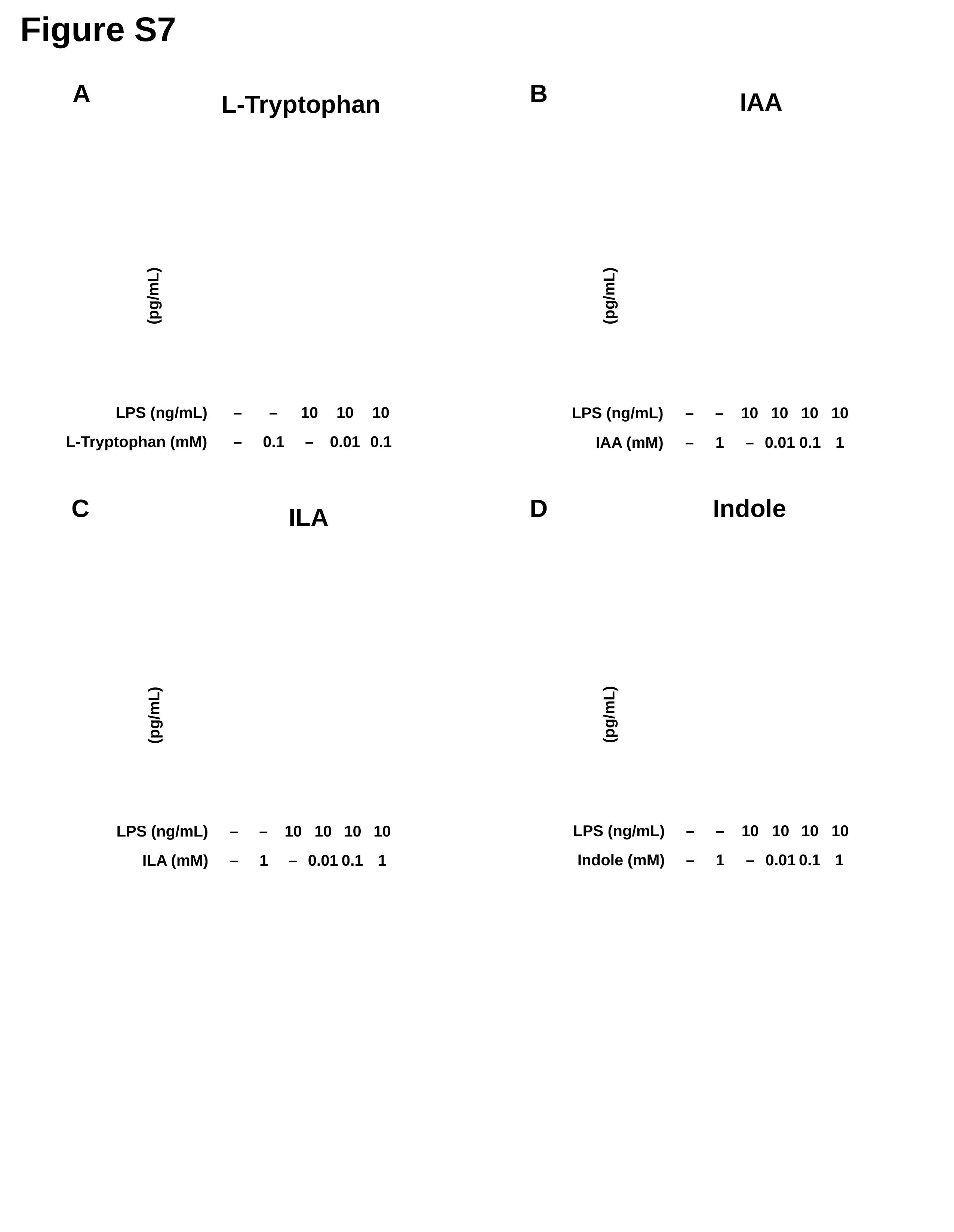

Figure S7
B
A
IAA
(pg/mL)
LPS (ng/mL)
–
–
10
10
10
10
IAA (mM)
–
1
–
0.01
0.1
1
L-Tryptophan
(pg/mL)
LPS (ng/mL)
–
–
10
10
10
L-Tryptophan (mM)
–
0.1
–
0.01
0.1
Indole
(pg/mL)
LPS (ng/mL)
–
–
10
10
10
10
Indole (mM)
–
1
–
0.01
0.1
1
C
D
ILA
(pg/mL)
10
LPS (ng/mL)
–
–
10
10
10
1
ILA (mM)
–
1
–
0.01
0.1

### Slide 8
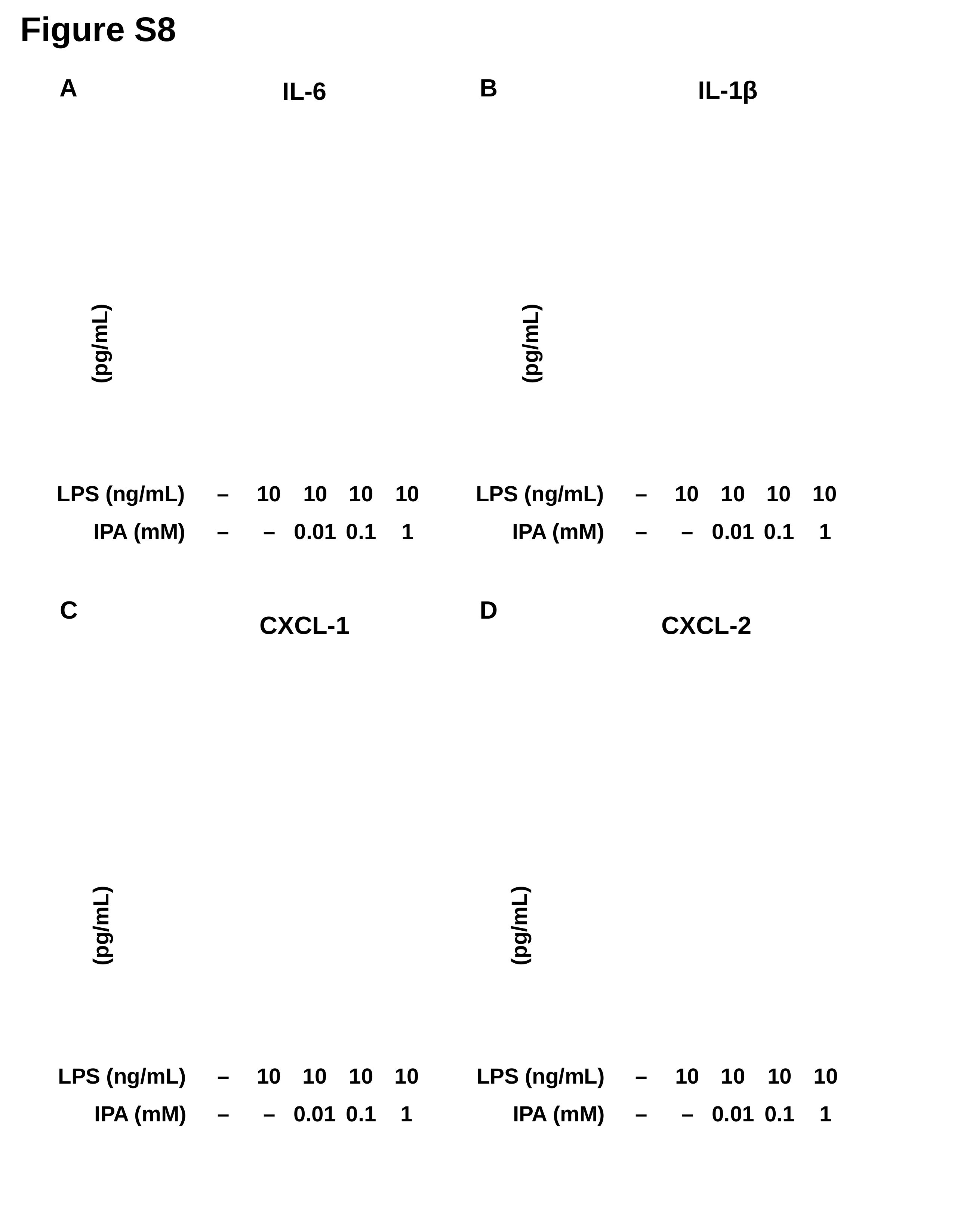

Figure S8
B
A
IL-1β
(pg/mL)
LPS (ng/mL)
–
10
10
10
10
IPA (mM)
–
–
0.01
0.1
1
IL-6
(pg/mL)
LPS (ng/mL)
–
10
10
10
10
IPA (mM)
–
–
0.01
0.1
1
D
C
CXCL-2
(pg/mL)
LPS (ng/mL)
–
10
10
10
10
IPA (mM)
–
–
0.01
0.1
1
CXCL-1
(pg/mL)
LPS (ng/mL)
–
10
10
10
10
IPA (mM)
–
–
0.01
0.1
1

### Slide 9
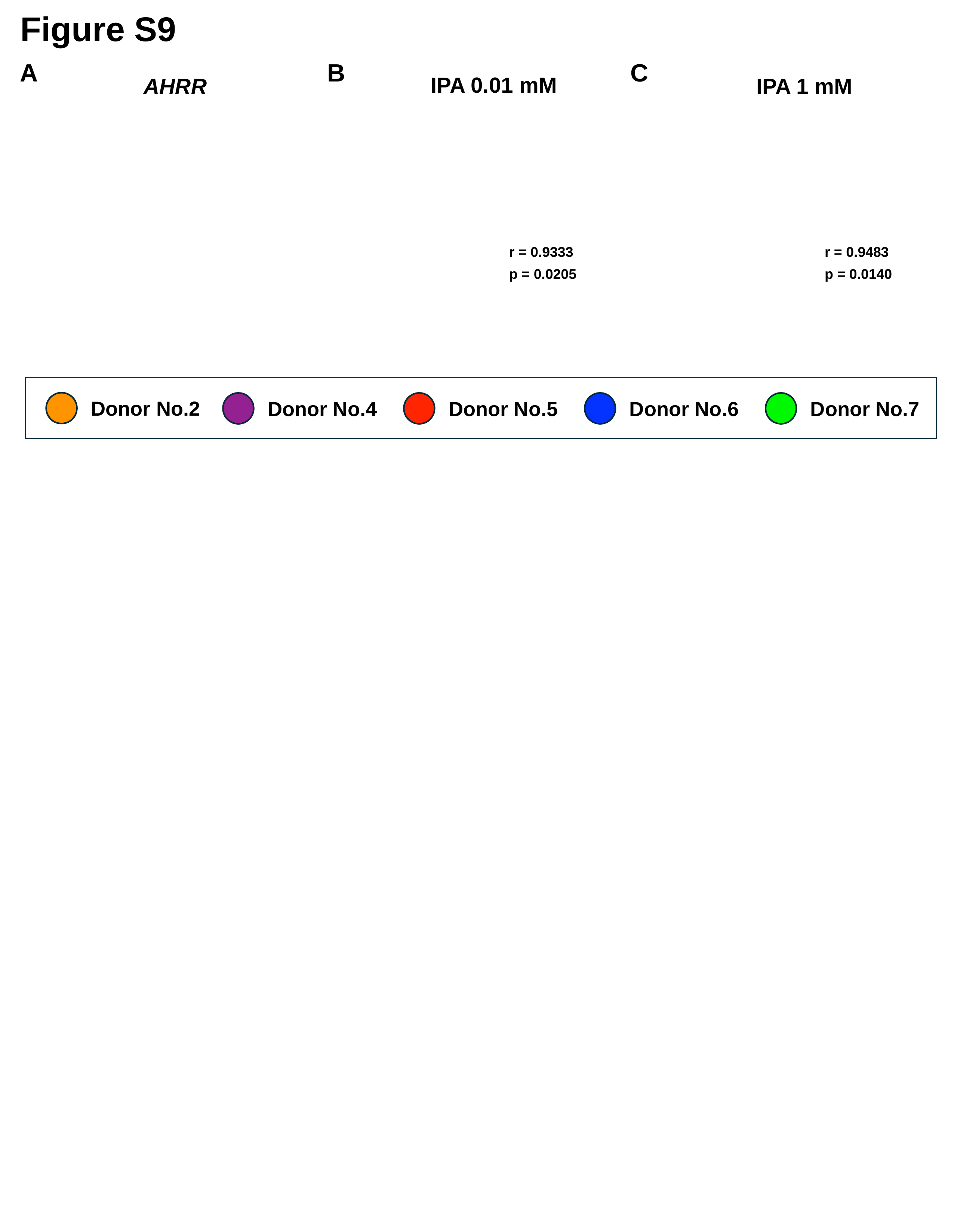

Figure S9
A
B
C
IPA 0.01 mM
AHRR
IPA 1 mM
r = 0.9333
r = 0.9483
p = 0.0205
p = 0.0140
Donor No.2
Donor No.4
Donor No.5
Donor No.6
Donor No.7

### Slide 10
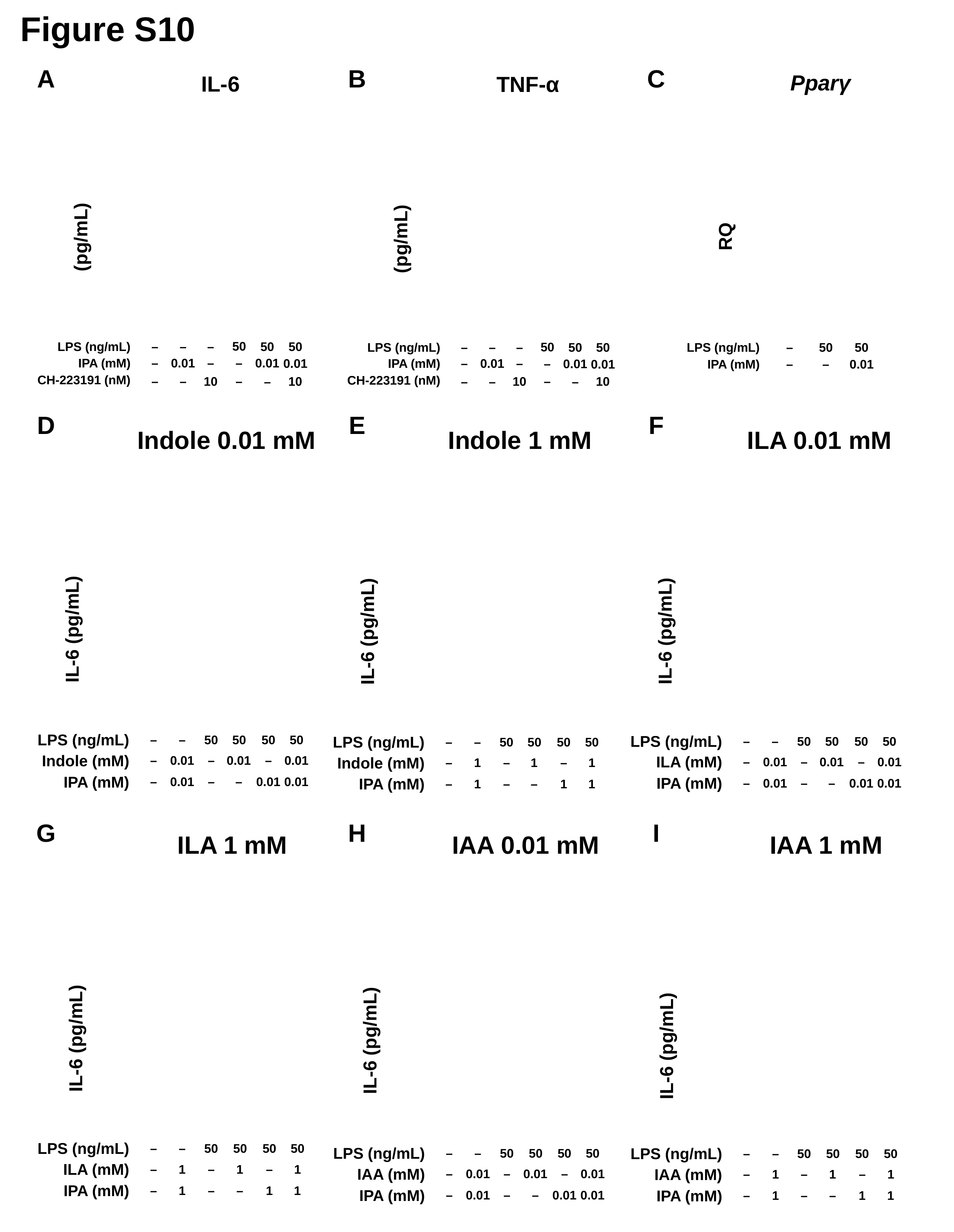

Figure S10
C
A
B
Pparγ
IL-6
TNF-α
RQ
(pg/mL)
(pg/mL)
50
–
50
50
LPS (ng/mL)
–
–
50
–
50
50
50
LPS (ng/mL)
–
–
LPS (ng/mL)
–
50
IPA (mM)
–
0.01
–
–
0.01
0.01
IPA (mM)
–
0.01
–
–
0.01
0.01
IPA (mM)
–
–
0.01
CH-223191 (nM)
CH-223191 (nM)
10
–
10
–
–
10
–
10
–
–
–
–
D
E
F
Indole 0.01 mM
IL-6 (pg/mL)
LPS (ng/mL)
–
–
50
50
50
50
Indole (mM)
–
0.01
–
0.01
–
0.01
IPA (mM)
–
0.01
–
–
0.01
0.01
Indole 1 mM
IL-6 (pg/mL)
LPS (ng/mL)
–
–
50
50
50
50
Indole (mM)
–
1
–
1
–
1
IPA (mM)
–
1
–
–
1
1
ILA 0.01 mM
IL-6 (pg/mL)
LPS (ng/mL)
–
–
50
50
50
50
ILA (mM)
–
0.01
–
0.01
–
0.01
IPA (mM)
–
0.01
–
–
0.01
0.01
G
H
I
IAA 1 mM
IL-6 (pg/mL)
LPS (ng/mL)
–
–
50
50
50
50
IAA (mM)
–
1
–
1
–
1
IPA (mM)
–
1
–
–
1
1
ILA 1 mM
IL-6 (pg/mL)
LPS (ng/mL)
–
–
50
50
50
50
ILA (mM)
–
1
–
1
–
1
IPA (mM)
–
1
–
–
1
1
IAA 0.01 mM
IL-6 (pg/mL)
LPS (ng/mL)
–
–
50
50
50
50
–
0.01
–
0.01
–
0.01
–
0.01
–
–
0.01
0.01
IAA (mM)
IPA (mM)
