## Supplemental Methods for "Dietary tryptophan mitigates lung ischemia-reperfusion injury via microbiota-derived indole-3-propionate and aryl hydrocarbon receptor signaling"

**Supplemental Materials and Methods**

**Animal care**

All animal experimentation protocols received approval from the Institutional Animal Care and Use Committee at the University of California San Francisco (protocol AN197325-01). Both male and female wild-type C57BL/6 mice, aged 10-15 weeks, were obtained from breeding colonies maintained systematically at UCSF. The mice were housed under strictly controlled environmental conditions, with a temperature of 21 ± 2°C, a relative humidity of 60 ± 5%, and a 12-hour light:12-hour dark cycle.

**Dietary intervention**

All murine subjects were maintained on a standard laboratory diet consisting of 0.24% tryptophan until the commencement of the dietary intervention. For the lung IR experiment, the subjects were randomly assigned to receive either a tryptophan-standard diet (Trp-Std, 0.18% tryptophan) or a tryptophan-rich diet (Trp-Rich, 0.60% tryptophan) for 14 days (Trp-Std, male: n =14, female: n = 13, Trp-Rich, male: n = 12, female: n = 11). For the intratracheal LPS challenge experiment, male mice were randomly allocated to the Trp-Std (n = 10) or Trp-Rich (n = 8) group. The diets were procured from Inotiv (Indianapolis, IN) and were specifically formulated to be isocaloric and isonitrogenous. To mitigate diet-induced alterations in gut microbiota composition attributed to fiber content, cellulose was standardized to 3% across both dietary formulations [1]. Food and water were administered *ad libitum* throughout the intervention. Comprehensive dietary compositions are delineated in Table S1.

**Left lung IR surgery**

A murine model of unilateral left pulmonary artery occlusion was established according to previously described methodologies [2-4]. In brief, the mice were anesthetized via intraperitoneal injection of 2,2,2-tribromoethanol (Avertin, Sigma-Aldrich, catalog number: T48402), followed by orotracheal intubation. Subsequently, buprenorphine was administered intramuscularly for analgesia (Covetrus North America, catalog number: 059122), and the mice were mechanically ventilated with a tidal volume of 225 µL, equivalent to 7.5 mL per kg for a 30 g mouse, at a respiratory rate of 180 breaths per minute. A left thoracotomy was executed through the second intercostal space, specifically between the second and third ribs. The left pulmonary artery was located and occluded utilizing a slipknot fashioned from 7-0 monofilament suture (Ethicon, Somerville, NJ, catalog number: 8696G). The tail of the suture was exteriorized to the anterior chest wall via a 27-gauge needle. Prior to chest closure, the left lung was re-expanded using sustained positive-pressure inflation. A topical local anesthetic was administered with 3 to 4 drops of 0.25% bupivacaine (Hospira, Lake Forest, IL, catalog number: NDC 0409-1159-19) before skin closure. The surgical procedure lasted approximately 25 to 30 minutes. Ischemia was sustained for a duration of 60 min, after which the ligature was released to commence reperfusion. Mice were euthanized one hour after reperfusion, and samples were collected. All mice underwent a comparable duration of mechanical ventilation, estimated at approximately 40-45 min. Post-extubation, the animals maintained spontaneous respiration throughout their recovery from anesthesia, during the entire ischemic interval, and throughout the reperfusion phase. Perioperative care adhered to institutional guidelines, and reporting complied with ARRIVE recommendations. The overall surgical success, operationally defined as survival to the predetermined endpoint, was approximately 85-90%. Animals that did not survive the IR procedure or the reperfusion phase due to technical complications, predominantly arising from injury to the left main bronchus or left pulmonary artery, were excluded from subsequent analysis. One Trp-Rich male cage (n = 5) was excluded because cage-mate fighting caused injuries. In addition, mice were excluded due to perioperative death attributable to technical complications from the microsurgical procedure (e.g. PA injury) (this last part can even be excluded altogether) (male Trp, n = 2; female Trp-Std, n = 2; male Trp-Rich, n = 1; female Trp-Rich, n = 2). Of note, body weight did not differ between groups (Figure S1A), and gross intestinal appearance was comparable (Figure S1B).

**Intratracheal lipopolysaccharide challenge**

Mice were anesthetized with 3% isoflurane by inhalation, and tracheally intubated. Lipopolysaccharide (LPS, *Escherichia coli* 0111:B4, Sigma-Aldrich, St. Louis, MO, catalog number: L4391) was administered intratracheally at 5 mg/kg [5,6]. One mouse in the Trp-Std group was excluded from analysis due to unsuccessful tracheal intubation and intratracheal LPS injection. Mice were mechanically ventilated for 30 s and then returned to their cages, where they were closely monitored until they fully recovered from anesthesia. Animals were subsequently monitored at 1-h intervals, and blood and lung samples were collected 6 h after LSP administration.

**Oxygen saturation monitoring and sample collection**

Before sample collection, peripheral oxygen saturation and respiratory rate were assessed using the MouseOx® Plus system (Starr Life Science Corp., Oakmont, PA). Oxygen saturation and respiratory rate were assessed in male mice only because the MouseOx neck-collar sensor is recommended for animals >25 g, and its use in smaller female mice could potentially compromise breathing. Furthermore, in LPS IT experiment, oxygen saturation was not assessed because of their moving. Following the physiological evaluation, the mice were anesthetized with 2,2,2-tribromoethanol, after which blood was collected from the inferior vena cava and the portal vein (PV) using heparinized syringes equipped with a 28-gauge needle. The collected samples were centrifuged at 14,000 g for 5 min, separating plasma, which was then snap-frozen in liquid nitrogen and stored at -80°C [7]. The caudal segment of the left lung was surgically excised and partitioned into two distinct aliquots. One aliquot was immersed in TRIzol reagent (Invitrogen Carlsbad, CA, catalog number: 15596018) for RNA extraction, while the other was preserved at -80°C for future homogenization and enzyme-linked immunosorbent assay (ELISA). ELISA quantified cytokine concentrations in lung homogenates, and gene expression was analyzed in lung tissue by reverse transcription quantitative polymerase chain reaction (RT-qPCR). The right lung was designated for the quantification of tryptophan metabolites utilizing liquid chromatography-mass spectrometry (LC-MS). Fecal pellets were collected from the descending colon and stored at -80°C for subsequent 16S rRNA gene sequencing and targeted measurements of tryptophan metabolites.

**Reagents and Cell Lines**

IPA (catalog number: 57400), ILA (catalog number: I5508), IAA (catalog number: I3750), indole (catalog number: I3408), L-tryptophan (catalog number: T8941), lipopolysaccharide (LPS) derived from *Escherichia coli* 0111:B4 (catalog number: L4391), and CH-223191 (catalog number: C8124) were purchased from Sigma-Aldrich (St. Louis, MO). Additionally, 2,3,7,8-tetrachlorodibenzo-p-dioxin (TCDD, catalog number: RPE-023S-1) was acquired from Neta Scientific (Hainesport, NJ). The MH-S cell line, representing a wild-type BALB/c mouse AM (catalog number: CRL-2019), was sourced from ATCC (Manassas, VA).

**Collection of Human Primary AMs by Bronchoalveolar Lavage**

Primary human AMs were procured from the lungs of adult donors who were unsuitable for transplantation. This lung was generously provided for research purposes by Dr. Michael Matthay at the University of California San Francisco. A bronchoalveolar lavage procedure was conducted by instilling ice-cold Dulbecco’s phosphate-buffered saline (PBS; Thermo Fisher Scientific, Waltham, MA, catalog number: 14190-144), enriched with 2 mM EDTA (Corning Costar, Corning, NY, catalog number: 46-034-CI) into the left upper lobe bronchus and subsequently aspirating the fluid in 50 mL aliquots to achieve a cumulative volume of 500 mL. Erythrocytes were eliminated utilizing 1x RBC Lysis Buffer (Thermo Fisher Scientific, catalog number: 00-4333-57) in accordance with the manufacturer’s guidelines. The cellular pellet was obtained by centrifugation at 300 g for 4 min at a temperature of 4°C, followed by resuspension in RPMI 1640 medium (Thermo Fisher Scientific, catalog: 72400-047) augmented with 10% fetal bovine serum (FBS, Thermo Fisher Scientific, catalog number: A5256801) and 1% penicillin-streptomycin (P/S, ThermoFisher Scientific, catalog number: 10378016). The proportion of AMs in the recovered cell population was confirmed by their characteristic autofluorescence using a fluorescence microscope (BZ-X1000, Keyence, Osaka, Japan). For the purpose of LPS stimulation assays, AMs were cultured in 48-well plates at a cellular density of 50,000 cells per well for ELISA and 12-well plates at a density of 160,000 cells per well for RT-qPCR and were allowed to adhere for a duration of 2 hours at 37°C within a humidified incubator containing 5% CO_2_. In an *ex vivo* nutritional IR experiment, 200,000 AMs per well were plated in a 48-well plate and incubated for 2 hours under identical conditions before subsequent treatments were applied.

***Ex vivo* nutritional IR in Human Primary AMs**

Following the seeding process, human primary AMs were co-treated overnight with LPS (200 ng/mL) and IPA at 0.01, 0.1, or 1 mM, in a temperature-controlled environment at 37°C in a humidified incubator supplemented with 5% CO_2_ [1,3]. Subsequently, the cells underwent two PBS washes and were incubated in PBS for 1 hour to simulate the ischemic phase, after which they were transferred to RPMI medium to simulate the reperfusion phase. Upon completion of this nutritional IR protocol, culture supernatants were harvested for quantification of IL-1β by ELISA. Following the removal of the supernatant, 200 µL of RPMI medium was added, and cell number and confluency were quantified using a Cytation 5 imaging reader (Agilent, Santa Clara, CA).

***Ex vivo* LPS Stimulation and IPA Treatment in Human Primary AMs**

Following a 2-hour incubation period post-seeding, human primary AMs were subjected to concurrent treatment with LPS (10 ng/mL) and IPA at specified concentrations (0.01, 0.1, or 1 mM) within a controlled environment maintained at 37°C in a humidified incubator enriched with 5% CO_2_. Subsequent to an overnight incubation period, the culture supernatants were harvested, and the concentrations of interleukin-6 (IL-6), interleukin-1 beta (IL-1β), chemokine (C-X-C motif) ligand 1 (CXCL-1), and chemokine (C-X-C motif) ligand 2 (CXCL-2) were measured utilizing ELISA methodology.

**Collection of Murine Primary AMs by Bronchoalveolar Lavage**

Primary murine AMs were collected by bronchoalveolar lavage after 2 weeks of dietary intervention in mice that did not undergo surgery. After surgical exposure of the trachea, a 20-gauge catheter was inserted, and prewarmed PBS (Thermo Fisher Scientific) containing 2 mM EDTA (Corning Costar) was instilled and gently aspirated in 1 mL aliquots to a total lavage volume of 10 mL. Cells were then pelleted by centrifugation at 300 × g for 4 min at 4°C, resuspended in RPMI 1640 medium supplemented with 10% FBS and 1% P/S, and plated in 48-well plates. For each individual mouse, recovered primary AMs were split into paired no-IR and IR conditions within the same diet group. Cells from Trp-Std mice were seeded into Trp-Std no-IR and Trp-Std IR wells, whereas cells from Trp-Rich mice were seeded into Trp-Rich no-IR and Trp-Rich IR wells, at 200,000 cells per well. Cells were allowed to adhere for 2 h at 37°C in a humidified incubator with 5% CO_2_ before *ex vivo* nutritional IR experiments.

***Ex vivo* nutritional IR in Murine Primary AMs**

After the adhesion period, murine primary AMs were exposed to LPS (200 ng/mL) overnight at 37°C in a humidified 5% CO_2_ incubator. Cells were then washed twice with PBS and incubated in PBS for 1 h to model the ischemic phase, followed by replacement with RPMI medium for 1 h to model reperfusion. At the end of the protocol, culture supernatants were collected, and IL-1β and IL-6 concentrations were measured by ELISA [1,3].

***In vitro* LPS stimulation using MH-S cells**

MH-S (50,000 cells per well) were cultured in 48-well plates utilizing RPMI 1640 medium enriched with 10% FBS and 1% P/S. The cells were maintained at 37°C in a humidified incubator with 5% CO_2_ for 24 hours. For the purpose of metabolite evaluation, cells were subjected to a pretreatment with IPA, ILA, IAA, indole, or L-tryptophan at specified concentrations (0.01, 0.1, or 1 mM) for a duration of 24 hours, followed by an overnight co-treatment with LPS at a concentration of 50 ng/mL, while maintaining the presence of the respective metabolite. After the incubation period, culture supernatants were harvested for ELISA analysis, and cells were collected for RT-qPCR. In an experiment using CH-223191, the cells were cultured overnight and subsequently subjected to a 4-hour pretreatment with CH-223191, followed by a 24-hour pretreatment with either IPA or TCDD, and a co-treatment with IPA or TCDD and LPS for an overnight duration.

**Sandwich Enzyme-Linked Immunosorbent Assay (ELISA)**

The concentrations of cytokines and chemokines in cell culture supernatants and mouse lung tissue were assessed using either mouse or human DuoSet ELISA kits (R&D Systems, Minneapolis, MN) according to the manufacturer's guidelines. Standard curves were established for each assay and used to determine analyte concentrations in the experimental samples.

**RT-qPCR**

RT-qPCR was performed using TaqMan inventoried gene expression assays to measure mRNA levels in pulmonary tissue and cultured cell lines. The assays encompassed *Actb* (Mm02619580_g1), *Gapdh* (Mm99999915_g1), *Rpl19* (Mm01606037_g1), *Pparg* (Mm00440940_m1), *Cyp1a1* (Mm00487218_m1), *Cyp1b1* (Mm00487229_m1), *ACTB* (Hs01060665_g1), *RPL32* (Hs00851655_g1), and *AHRR* (Hs01005075_m1) (all sourced from Thermo Fisher Scientific). The left lung tissue was homogenized using a Tissue-Tearor homogenizer (BioSpec Products, Bartlesville, OK). Total RNA was extracted employing TRIzol reagent and subsequently purified utilizing the RNeasy Mini Kit (Qiagen, Venlo, Netherlands, catalog number: 74104). cDNA was synthesized from 1 µg of total RNA by means of the High-Capacity RNA-to-cDNA Kit (Applied Biosystems, Foster City, CA, catalog number: 4387406). qPCR reactions were prepared using TaqMan Fast Advanced Master Mix (Applied Biosystems, catalog number: 444557) in conjunction with the designated TaqMan Gene Expression Assays and were executed on QuantStudio 6/7 Flex Real-Time PCR Systems (Applied Biosystems). Cycling parameters consisted of an initial incubation at 95°C for 20 s, followed by 40 cycles of denaturation at 95°C for 1 s and annealing/extension at 60°C for 20 s. Each sample was analyzed in technical triplicate, and the mean Ct was used for subsequent analysis. The mean Ct values of *Actb*, *Gapdh*, and *Rpl19* served as internal reference genes for calculating ΔCt. Relative expression levels were computed utilizing the 2^-ΔΔCt method and reported as relative quantification [8,9].

**16S rRNA gene sequencing**

For microbiome analysis, we analyzed the first four mice enrolled per group (n = 4 per group). Genomic DNA was extracted from previously collected fecal samples using the QIAamp PowerFecal Pro DNA Kit (QIAGEN, catalog number: 51804) according to the manufacturer’s instructions. Briefly, approximately 25 mg of fecal material was processed in a biosafety cabinet and transferred into PowerBead tubes containing the kit lysis buffer (C1). Mechanical disruption was performed using a TissueLyser II (QIAGEN, catalog number: 9003240) at 25 Hz for 5 min with 2 times to ensure efficient lysis. For the extraction negative control, 500 µL nuclease-free water was added to a PowerBead tube containing C1 buffer and processed in parallel through all subsequent steps. DNA was eluted in 75 µL nuclease-free water and stored at -20°C until downstream analyses. DNA quantity and purity were assessed using a NanoDrop spectrophotometer (Thermo Fisher Scientific) by measuring absorbance ratios at A260/A280 and A260/A230. The V4 region of the bacterial 16S rRNA gene was amplified by PCR using the 515F/806R primer pair. Reverse primers included sample-specific barcode sequences and Illumina adapter sequences to enable multiplexing and flow-cell binding. Amplification was performed in 25 µL reactions containing Hot Start DNA polymerase (Hot Start Ex Taq, Takara Bio, Shiga, Japan), 1× reaction buffer, 0.4 µM of each primer, bovine serum albumin (BSA), dNTPs, and 10 ng of template DNA. PCR conditions were 98°C for 2 min, followed by 30 cycles of 98°C for 20 s, 50°C for 30 s, and 72°C for 45 s, with a final extension at 72°C for 10 min. Amplicons were quantified using a Qubit dsDNA HS assay and verified by agarose gel electrophoresis. Libraries were pooled at equimolar concentrations using an automated liquid-handling platform, purified with AMPure SPRI beads, and assessed for size distribution and concentration using a Bioanalyzer and Qubit. The pooled library was diluted to 2 nM, spiked with 10% PhiX, and sequenced on an Illumina MiSeq system (2×300 bp). Raw base-call files were converted to FASTQ format (bcl2fastq v2.20), and reads were demultiplexed by barcode prior to downstream analyses. Demultiplexed paired-end reads were subsequently processed using a DADA2-based workflow in MicrobiomeAnalyst, and taxonomy was assigned against SILVA v138, as described below.

**Quantification of tryptophan metabolites**

Tryptophan-related metabolites were extracted from mouse feces, PV plasma, and lung using matrix-specific protocols with isotope-labeled internal standards. For PV plasma, 30 µL was mixed with 120 µL of extract solvent (1:1:1 methanol:acetonitrile:acetone) containing internal standards, vortexed, incubated on ice for 10 min, and centrifuged (15,000 x g, 10 min). Supernatants (100 µL) were transferred to autosampler vials and dried under nitrogen. For feces and lung, weighed samples were extracted with 1:1:1:1 methanol:acetonitrile:water:acetone containing internal standards (16 µL/mg tissue) in Precellys tubes with a 2.7-mm stainless steel bead, homogenized (6200 rpm, 3 cycles of 20 s shake/30 s rest at 4°C), and centrifuged (14,000 rpm, 10 min, 4°C). Supernatants (150 µL) were dried under nitrogen. Dried extracts were reconstituted in 80:20 water/methanol (50 µL for plasma; 75 µL for lung and feces). A pooled quality-control sample (aliquots combined from all extracts) was processed identically and injected periodically throughout the run. Quantitation used a 7-point calibration curve prepared alongside samples. The internal standard mix included Tryptophan-D5, Kynurenic Acid-D5, L-Kynurenine-D4, 5-Hydroxyanthranilic Acid-D3; standard ranges were 0.5-40 µM for L-tryptophan, 5-hydroxytryptophan, and 5-hydroxyindolacetic acid, and 25 nM-2 µM for 3-hydroxyanthranilic acid, kynurenic acid, L-kynurenine, and related metabolites. LC-MS analysis was performed on an Agilent 1290 Infinity II coupled to an Agilent 6545 qTOF with JetStream ESI, using an Acquity HSS T3 column (2.1x100 mm, 1.7 µm) at 55°C with a flow rate of 0.45 mL/min and mobile phases of water (A), methanol (B), and 2.5% formic acid (C). The 17.1-min gradient was 0% B/4% C (0 min), 99% B/1% C (10-17 min), and 0% B/1% C (17.1 min), followed by 3 min reconditioning. Samples were acquired in positive and negative ion modes (10 µL injection) with drying gas at 275°C (12 L/min), nebulizer at 45 psig, sheath gas at 325°C (12 L/min), capillary voltage at 4000 V, and internal reference mass correction. Raw data were processed in MassHunter Quantitative Analysis (B.10.00), normalized to the nearest isotope-labeled internal standard, and quantified against the linear calibration curve; when detected in both ion modes, results were reported from the mode with the higher signal and lower pooled-QC CV. Tissue concentrations were normalized to input mass and reported as µM/mg tissue.

**Multi-omics integration and correlation analyses**

Fecal samples from male mice were collected after a 2-week dietary intervention (Trp-Std vs Trp-Rich; n = 4 per group). Paired-end FASTQ files were processed on the MicrobiomeAnalyst (microbiomeanalyst.ca) [10-12]. Raw reads were denoised with a DADA2-based pipeline: forward/reverse truncation at 259/219 bp, TrimLeft/TrimRight = 10/10, maxEE = 2/2, maxN = 0, minQ = 1, truncQ = 2; chimeras were removed, and amplicon sequence variants (ASVs) were inferred. Taxonomy was assigned with SILVA v138. For Marker Data Profiling, features were filtered with a low-count threshold (minimum count = 4), prevalence (≥20% of samples), and low-variance removal (bottom 10% by inter-quartile range). For β-diversity, we uploaded a phylogenetic tree built from ASV representative sequences in R using DECIPHER (MSA) and phangorn GTR (midpoint-rooted), and computed PCoA on Bray-Curtis, unweighted UniFrac, and weighted UniFrac distances in the MicrobiomeAnalyst. Group separation was tested using PERMANOVA. For community summaries and correlations, count tables were total-sum scaled (TSS) and centered log-ratio (CLR) transformed. Associations between genus/species abundances and lung cytokines and AhR signaling readouts (IL-1β, IL-6, *Cyp1a1*, *Cyp1b1*) were assessed using Spearman correlations. Differential abundance between diets was assessed at the genus and species levels using DESeq2 on raw counts (internal size-factor normalization), and false discovery rate-adjusted p-values are reported. Predicted functional potential from 16S was obtained with Tax4Fun2 (SILVA) to generate KEGG Orthology (KO) profiles. Single-factor comparisons of KO tables between diets were conducted with metagenomeSeq (fitFeature), and pathway-level (KEGG LE) summaries were derived in MicrobiomeAnalyst. All thresholds (e.g., adjusted p < 0.05) and figure options (e.g., box/violin plots, PCoA with 95% ellipses) match these settings unless otherwise indicated.

Targeted tryptophan metabolite concentration data from feces, PV plasma, and lung were analyzed in MetaboAnalyst 6.0 using the Statistical Analysis one-factor module with samples in rows [13]. Data were filtered using default variance and abundance options, and missing values were imputed using a left-censored approach with LoD set to one-fifth of the minimum positive value. Data were log10-transformed, and Preto scaled before analyses. Group separation was explored using PCoA and PLS-DA with three components. For univariate comparisons between Trp-Std and Trp-Rich groups, non-parametric t-tests with unequal variance option were applied with FDR correction, and volcano plots were generated using an FDR-adjusted p-value threshold of 0.05 and a fold change threshold of 1.5 with the comparison defined as Trp-Rich over Trp-Std. To relate metabolite patterns to external readouts, the Statistical Analysis metadata table module was used. Spearman rank correlations were computed between selected metadata targets and metabolite features. For visualization, hierarchical clustering heatmaps were generated from normalized data with feature autoscaling, Pearson distance, and complete linkage clustering, with sample ordering by group and annotation tracks including Group, Lung IL-1β, and *Cyp1a1*.

**Statistical Analyses**

Data were presented as mean ± SD. Data from *in vitro* studies comparing two conditions were analyzed using an unpaired t-test or a Mann-Whitney test to generate p-values, after confirming normality with the Shapiro-Wilk test. For comparisons involving more than two conditions, we used one-way ANOVA, as appropriate, with Tukey’s post hoc test. In *ex vivo* experiments using human AMs, paired t-tests, Wilcoxon signed-rank tests, or repeated-measures one-way ANOVA followed by Tukey’s post hoc tests were performed. GraphPad Prism was used for statistical analysis (GraphPad Software ver.10, La Jolla, CA). For microbiome-host correlation analyses (Spearman), given the small sample size (n=4 per group), results were considered exploratory and unadjusted p-values are reported without false discovery rate correction. Unadjusted or adjusted p-values < 0.05 were considered significant. P*-*values are presented in the figures as follows: *< 0.05; **< 0.01; ***< 0.001; ****< 0.0001.
