## Supplemental Tables for "Dietary tryptophan mitigates lung ischemia-reperfusion injury via microbiota-derived indole-3-propionate and aryl hydrocarbon receptor signaling"

Table S1. Formulas of tryptophan-standard (Trp-Std) and tryptophan-rich (Trp-Rich) diets.

|  | Trp-Std | Trp-Rich |
| --- | --- | --- |
| Nutrient information | | |
| Protein (% by weight) | 15.4 | 15.4 |
| Protein (% kcal) | 15.6 | 15.6 |
| Carbohydrate (% by weight) | 64.9 | 64.9 |
| Carbohydrate (% kcal) | 66.0 | 66.0 |
| Fat (% by weight) | 8.0 | 8.0 |
| Fat (% kcal) | 18.3 | 18.3 |
| Kcal/g | 3.9 | 3.9 |
| Formula (g/Kg) | | |
| Sucrose | 344.98 | 345.073 |
| Corn Starch | 150.0 | 150.0 |
| Maltodextrin | 150.0 | 150.0 |
| Soybean Oil | 80.0 | 80.0 |
| Cellulose | 30.0 | 30.0 |
| Mineral Mix, AIN-93M-MX (94049) | 35.0 | 35.0 |
| Calcium Phosphate, monobasic, monohydrate | 8.2 | 8.2 |
| Vitamin Mix, AIN-93-VX (94047) | 19.5 | 19.5 |
| Choline Bitartrate | 2.7 | 2.7 |
| Tert-Butylhydroquinone, antioxidant | 0.02 | 0.02 |
| Amino Acids | | |
| L-Alanine | 3.5 | 2.976 |
| L-Arginine HCl | 12.1 | 12.1 |
| L-Asparagine | 6.0 | 5.612 |
| L-Aspartic Acid | 3.5 | 2.718 |
| L-Cystine | 3.5 | 3.5 |
| L-Glutamic Acid | 40.0 | 39.135 |
| Glycine | 23.3 | 22.859 |
| L-Histidine HCl, monohydrate | 4.5 | 4.5 |
| L-Isoleucine | 8.2 | 8.2 |
| L-Leucine | 11.1 | 11.1 |
| L-Lysine HCl | 18.0 | 18.0 |
| L-Methionine | 8.2 | 8.2 |
| L-Phenylalanine | 7.5 | 7.5 |
| L-Proline | 3.5 | 2.824 |
| L-Serine | 3.5 | 2.883 |
| L-Threonine | 8.2 | 8.2 |
| L-Tryptophan | 1.8 | 6.0 |
| L-Tyrosine | 5.0 | 5.0 |
| L-Valine | 8.2 | 8.2 |

Table S2. Characteristics of lung donors.

| Donor No. | Sex | Age  (yrs) | Race | Height (cm) | Weight  (cm) | BMI  (kg/m^2^) | Medical history | Smoking history |
| --- | --- | --- | --- | --- | --- | --- | --- | --- |
| 1 | Female | 54 | White | 154 | 63.0 | 26.6 | - Hypertension - Heavy alcohol use | No |
| 2 | Female | 53 | White | 163 | 89.8 | 33.8 | - Hypertension | Yes |
| 3 | Female | 61 | Black | 159 | 75.0 | 29.7 | - Hypertension - Diabetes mellites | No |
| 4 | Male | 50 | Asian | 171 | 87.5 | 29.9 | - Hypertension - Myocardial infarction | No |
| 5 | Male | 60 | Asian | 164 | 83.3 | 31.0 | - Heavy alcohol use | No |
| 6 | Male | 36 | White | 178 | 102.3 | 32.3 | - Hypertension - Heavy alcohol use | No |
| 7 | Male | 53 | White | 180 | 110.4 | 34.1 | - Hypertension | No |

Table S3. Differential abundance analysis at the *Phylum* level.

| Taxa | Log_2_FC | SEM of Log_2_FC | Adjusted p value |
| --- | --- | --- | --- |
| *Firmicutes* | 1.3387 | 0.33641 | 0.00055274 |
| *Campylobacterota* | -2.0448 | 0.76059 | 0.020944 |
| *Cyanobacteria* | -4.3149 | 1.6232 | 0.020944 |
| *Actinobacteriota* | 1.9058 | 0.9413 | 0.085815 |
| *Verrucomicrobiota* | -0.80458 | 0.62856 | 0.3164 |
| *Proteobacteria* | -1.06258 | 1.3758 | 0.3164 |
| *Bacteroidota* | 0.071282 | 0.42389 | 0.9768 |
| *Desulfobacterota* | -0.019251 | 0.66192 | 0.9768 |

Table S4. Differential abundance analysis at the *Genus* level.

| Taxa | Log_2_FC | SEM of Log_2_FC | Adjusted p value |
| --- | --- | --- | --- |
| *Ileibacterium* | 16.978 | 1.2812 | 2.1115e-38 |
| *Dubosiella* | -15.382 | 1.2002 | 3.1698e-36 |
| *Rikenellaceae_RC9_gut_group* | -10.089 | 1.1183 | 2.9659e-18 |
| *Acetatifactor* | 10.393 | 1.2658 | 2.6368e-15 |
| *Staphylococcus* | -9.478 | 1.3181 | 6.1876e-12 |
| *Faecalibaculum* | 3.4805 | 0.53417 | 5.7897e-10 |
| *HT002* | 9.4749 | 2.0624 | 2.9786e-05 |
| *Jeotgalicoccus* | -7.2005 | 1.6214 | 5.3217e-05 |
| *Odoribacter* | 9.033 | 2.0448 | 5.3217e-05 |
| *Lactobacillus* | 2.7661 | 0.66833 | 0.00016757 |
| *Bifidobacterium* | 4.2892 | 1.123 | 0.00058372 |
| *Oscillibacter* | -1.5124 | 0.43342 | 0.0019352 |
| *Candidatus_Stoquefichus* | -5.6569 | 1.8125 | 0.0066533 |
| *Intestinimonas* | -1.4859 | 0.51882 | 0.014012 |
| *Helicobacter* | -2.382 | 0.83594 | 0.014012 |
| *Muribaculum* | 1.7746 | 0.67407 | 0.02541 |
| *UBA1819* | 6.4208 | 2.5173 | 0.030358 |
| *Lachnospiraceae_UCG_004* | -2.0827 | 0.89133 | 0.051886 |
| *Streptococcus* | -5.6307 | 2.5307 | 0.065903 |
| *Anaerotruncus* | -1.1149 | 0.52821 | 0.0835 |
| *Anaerostipes* | 5.6637 | 2.7099 | 0.083696 |
| *Desulfovibrio* | -3.1201 | 1.5988 | 0.11126 |
| *Marvinbryantia* | -1.7829 | 0.98156 | 0.14464 |
| *Coriobacteriaceae_UCG_002* | -2.5935 | 1.4618 | 0.1516 |
| *Escherichia_Shigella* | -5.0174 | 2.8561 | 0.1516 |
| *Not_Assigned* | -0.34148 | 0.25211 | 0.32416 |
| *Incertae_Sedis* | 0.62351 | 0.5057 | 0.38173 |
| *Parabacteroides* | -0.62703 | 0.52025 | 0.38173 |
| *Roseburia* | 1.288 | 1.0788 | 0.38173 |
| *Akkermansia* | -0.95202 | 0.8078 | 0.38173 |
| *Alistipes* | -0.40604 | 0.40908 | 0.49692 |
| *Ureaplasma* | 1.8694 | 2.0837 | 0.55447 |
| *Tuzzerella* | 0.4187 | 0.53624 | 0.62653 |
| *Erysipelatoclostridium* | 1.2795 | 1.6708 | 0.62653 |
| *Bacteroides* | -0.41373 | 0.62084 | 0.69278 |
| *Clostridium_sensu_stricto_1* | 0.62649 | 1.0269 | 0.72242 |
| *UCG_009* | 0.84421 | 1.5074 | 0.74162 |
| *Ligilactobacillus* | -0.33066 | 0.60894 | 0.74162 |
| *Turicibacter* | 0.48226 | 0.99721 | 0.77265 |

Table S4. (Continued)

| Taxa | Log_2_FC | SEM of Log_2_FC | Adjusted p value |
| --- | --- | --- | --- |
| *Colidextribacter* | 0.20721 | 0.44822 | 0.77265 |
| *Lachnospiraceae_NK4A136_group* | -0.21189 | 0.54097 | 0.80704 |
| *Lachnospiraceae_FCS020_group* | 0.18418 | 0.4885 | 0.80704 |
| *Bilophila* | -0.70848 | 2.0105 | 0.80879 |
| *Lachnoclostridium* | -0.089552 | 0.39375 | 0.89464 |
| *GCA_900066575* | -0.097943 | 0.51828 | 0.90678 |
| *A2* | -0.048655 | 0.35695 | 0.93034 |
| *Lachnospiraceae_UCG_006* | 0.070177 | 1.0798 | 0.96836 |
| *Enterorhabdus* | -0.013134 | 2.5065 | 0.99582 |

Table S5. Differential abundance analysis at the *Species* level.

| Taxa | Log_2_FC | SEM of Log_2_FC | Adjusted p value |
| --- | --- | --- | --- |
| *Ileibacterium_valens* | 16.909 | 1.2462 | 9.2334E-41 |
| *Faecalibaculum_rodentium* | 3.3688 | 0.51606 | 5.0007E-10 |
| *Lactobacillus_intestinalis* | 7.0342 | 2.0854 | 0.0037164 |
| *Helicobacter_ganmani* | -2.2717 | 0.73314 | 0.0072917 |
| *Muribaculum_intestinale* | 1.6368 | 0.61291 | 0.022722 |
| *Streptococcus_danieliae* | -5.7481 | 2.2937 | 0.030523 |
| *Not_Assigned* | -0.932 | 0.42789 | 0.055825 |
| *Anaerostipes_caccae* | 5.7241 | 2.6341 | 0.055825 |
| *Akkermansia_municiphila* | -1.1203 | 0.62685 | 0.11732 |
| *Bacteroides_acidifaciens* | -0.94694 | 0.53768 | 0.11732 |
| *Parabacteroides_distasonis* | -0.81528 | 0.51912 | 0.15343 |
| *Bacteroides_uniformis* | 1.7517 | 1.135 | 0.15343 |
| *Helicobacter_hepaticus* | -1.148 | 1.2938 | 0.43259 |
| *Lachnospiraceae_NK4A136_group_bacterium* | 0.268 | 0.6674 | 0.73715 |
| *Anaerotruncus_colihominis* | 0.3299 | 1.4515 | 0.82021 |
